# Optimal prediction with resource constraints using the information bottleneck

**DOI:** 10.1101/2020.04.29.069179

**Authors:** Vedant Sachdeva, Thierry Mora, Aleksandra M. Walczak, Stephanie Palmer

## Abstract

Responding to stimuli requires that organisms encode information about the external world. Not all parts of the signal are important for behavior, and resource limitations demand that signals be compressed. Prediction of the future input is widely beneficial in many biological systems. We compute the trade-offs between representing the past faithfully and predicting the future for input dynamics with different levels of complexity. For motion prediction, we show that, depending on the parameters in the input dynamics, velocity or position coordinates prove more predictive. We identify the properties of global, transferrable strategies for time-varying stimuli. For non-Markovian dynamics we explore the role of long-term memory of the internal representation. Lastly, we show that prediction in evolutionary population dynamics is linked to clustering allele frequencies into non-overlapping memories, revealing a very different prediction strategy from motion prediction.

## I. INTRODUCTION

How biological systems represent external stimuli is critical to their behavior. The efficient coding hypothesis, which states that neural systems extract as much information as possible from the external world, given basic capacity constraints, has been successful in explaining some early sensory representations in neuroscience. Barlow suggested sensory circuits may reduce redundancy in the neural code and minimize metabolic costs for signal transmission [1–4]. However, not all external stimuli are as important to an organism, and behavioral and environmental constraints need to be integrated into this picture to more broadly characterize biological encoding. Delays in signal transduction in biological systems mean that predicting external stimuli efficiently can confer benefits to biological systems [5–7], making prediction a general goal in biological sensing.

Evidence that representations constructed by sensory systems efficiently encode predictive information has been found in the visual and olfactory systems [8–10]. Molecular networks have also been shown to be predictive of future states, suggesting prediction may be one of the underlying principles of biological computation [11, 12]. However, the coding capacity of biological systems is limited because they cannot provide arbitrarily high precision about their inputs: limited metabolic resources and other sources of internal noise impose finite precision signal encoding. Given these trade-offs, one way to efficiently encode the history of an external stimulus is to keep only the information relevant for the prediction of the future input [12–14]. Here, we explore how optimal predictions might be encoded by neural and molecular systems using a variety of dynamical inputs that explore a range of temporal correlation structures. We solve the ‘information bottleneck’ problem in each of these scenarios and describe the optimal encoding structure in each case.

The information bottleneck framework allows us to define a ‘relevance’ variable in the encoded sensory stream, which we take to be the future behavior of that input. Solving the bottleneck problem allows us to optimally estimate the future state of the external stimulus, given a certain amount of information retained about the past. In general, prediction of the future coordinates of a system, *X*_*t*+∆*t*_ reduces to knowing the precise historical coordinates of the stimulus *X*_*t*_ and an exact knowledge of the temporal correlations in the system. These rules and temporal correlations can be thought of as arising from two parts: a deterministic portion, described by a function of the previous coordinates, 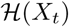, and the noise internal to the system, *ξ*(*t*). Knowing the actual realization of the noise *ξ*(*t*) reduces the prediction problem to simply integrating the stochastic equations of motion forward in time. If the exact realization of the noise if not known, we can still perform a stochastic prediction by calculating the future form of the probability distribution of the variable *X*_*t*_ or its moments [15, 16]. The higher-order moments yield an estimate of *X*_*t*_ and the uncertainty in the our estimate. However, biological systems cannot precisely know *X*_*t*_ due to inherently limited readout precision [17, 18] and limited availability of resources tasked with remembering the measured statistics.

Constructing internal representations of sensory stimuli illustrates a tension between predicting the future, for which the past must be known with higher certainty, and compression of knowledge of the past, due to finite resources. We explore this intrinsic trade-off using the information bottleneck (IB) approach proposed by Tishby et. al. [13]. This method assumes that the input variable, in our case the past signal *X*_*t*_, can be used to make inferences about the relevance variable, in our case the future signal *X*_*t*+∆*t*_. By introducing a representation variable, 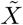, we can construct the conditional distribution of the representation variable on the input variable 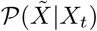 to be maximally informative of the output variable (Fig. 1).

**FIG. 1:**
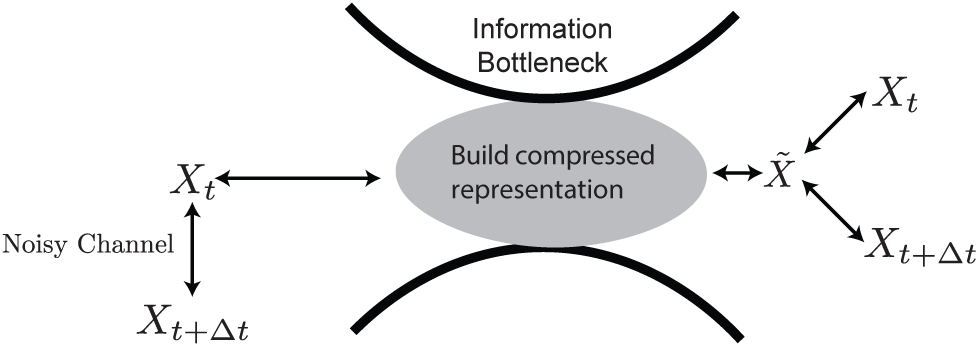
A schematic representation our predictive information bottleneck. On the left hand side, we have coordinates *X*_*t*_ evolving in time, subject to noise to give *X*_*t*+∆*t*_. We construct a representation, 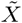, that compresses the past input (minimizes 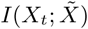) while retaining as much information about the future (maximizes 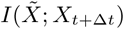) up to the weighting of the prediction compared to the compression set by *β*.

Formally, the representation is constructed by optimizing the objective function,

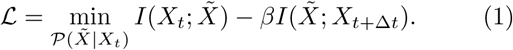

Each term is the mutual information between two variables: the first between the past input and estimate of the past given our representation model, 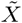, and the second between 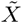 and future input. The tradeoff parameter, *β*, controls how much future information we want 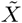 to retain as it is maximally compressed. For large *β*, the representation variable must be maximally informative about *X*_*t*+∆*t*_, and will have, in general, the lowest compression. Small *β* means less information is retained about the future and high, lossy compression is allowed.

The causal relationship between the past and the future results in a data processing inequality, 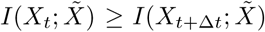, meaning that the information generated about the future cannot exceed the amount encoded about the past [19]. Additionally, the information about the past that the representation can extract is bounded by the amount of information the uncompressed past, itself, contains about the future, 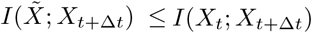.

We use this framework to study prediction in two well-studied dynamical systems with ties to biological data: the stochastically driven damped harmonic oscillator (SDDHO) and the Wright-Fisher model. We look simultaneously at these two different systems to gain intuition about how different types of dynamics influence the ability of a finite and noisy system to make accurate predictions. We further consider two types of SDDHO processes with different noise profiles to study the effect of noise correlations on prediction. Our exploration of the SDDHO system has a two-fold motivation: it is the simplest possible continuous stochastic system whose full dynamics can be solved exactly. Additionally, a visual stimulus in the form of a moving bar that was driven by an SDDHO process was used in retinal response studies [9, 20, 21]. The Wright-Fisher model [22] is a canonical model of evolution [23] for which has been used to consider how the adaptive immune system predicts the future state of the pathogenic environment [11, 24].

## II. RESULTS

### A. The Stochastically Driven Damped Harmonic Oscillator

Previous work explored the ability of the retina to construct an optimally predictive internal representation of a dynamic stimulus. Palmer et al [9] recorded the response of a salamander retina to a moving bar stimulus with SDDHO dynamics. In this case, the spike trains in the retina encode information about the past stimuli in a near-optimally predictive way [9]. In order for optimal prediction to be possible, the retina should encode the position and velocity as dictated by the information bottleneck solution to the problem, for the retina’s given level of compression of the visual input. Inspired by this experiment, we explore the optimal predictive encoding schemes as a function of the parameters in the dynamics, and we describe the optimal solution across the entire parameter space of the model, over a wide range of desired prediction timescales.

We consider the dynamics of a mass *m* in a viscous medium attached to a spring receiving noisy velocity kicks generated by a temporally uncorrelated Gaussian process, as depicted in Figure 2a. Equations of motion are introduced in terms of physical variables 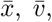, and 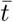 (bars will be dropped later when referring to rescaled variables), which evolve according to

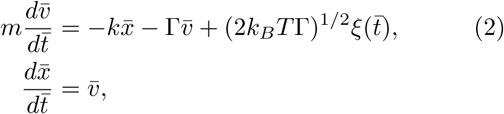

where *k* is the spring constant, Γ the damping parameter, *k*_*B*_ the Boltzmann constant, *T* temperature, 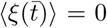, and 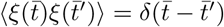. We rewrite the equation with

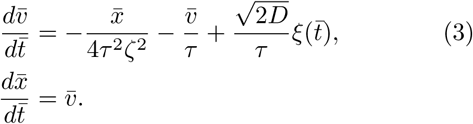

**FIG. 2:**
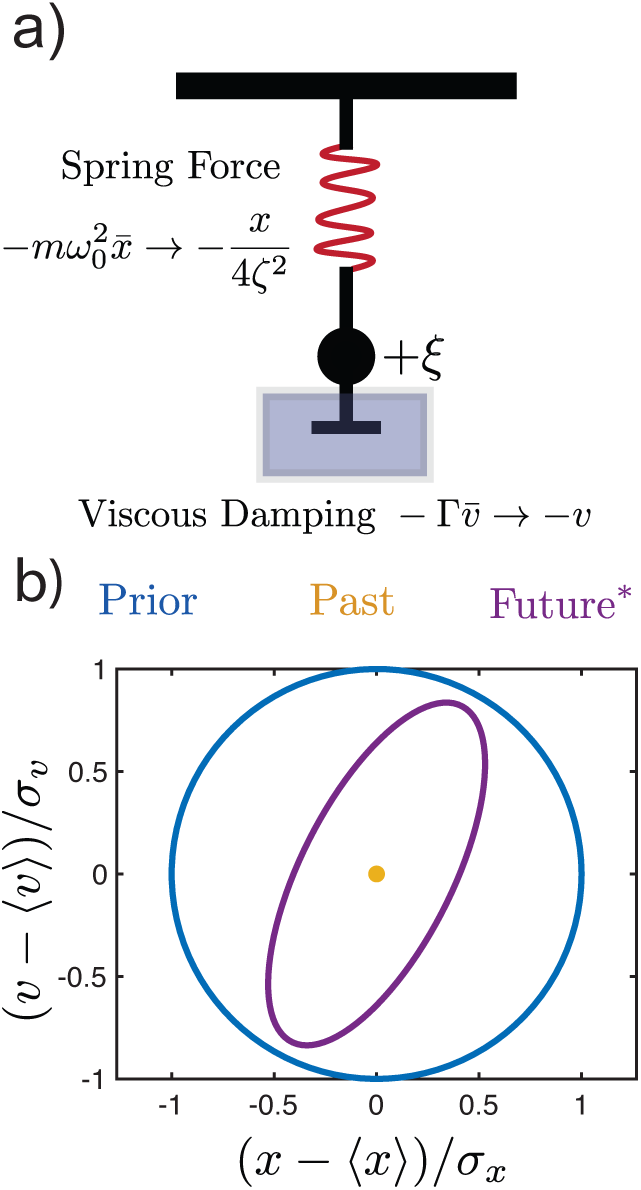
Schematic of the stochastically driven damped harmonic oscillator (SDDHO). (a) The SDDHO consists of a mass attached to a spring undergoing viscous damping and experiencing Gaussian thermal noise of magnitude. There are two parameters to be explored in this model: 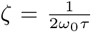 and 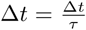. (b) We can represent the statistics of the stimulus through error ellipses. 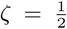, and ∆*t* = 1, we plot two-dimensional confidence intervals under various conditions. In blue, we plot the two-dimensional confidence interval of the prior. In yellow, we plot the certainty with which we measure the position and velocity at time *t*. Here, it is measured with infinite precision, meaning *I*_past_ → ∞. In purple, we plot the two-dimensional confidence interval of the future conditioned on the measurement given in yellow, for this particular choice of parameters. Precise knowledge of the past coordinates reduces the our uncertainty about the future position and velocity (as compared to the prior), as depicted by the smaller area of the purple ellipse.

We introduce a dimensionless parameter, the damping coefficient, *ζ* = 1/(2*ω*_0_*τ*). When *ζ* < 1, the motion of the mass will be oscillatory. When *ζ* ≥ 1, the motion will be non-oscillatory. Additionally, we note that the equipartition theorem tells us that 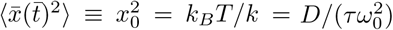

Expressing the equations of motion in terms of *ζ*, *τ*, and *x*_0_, we obtain

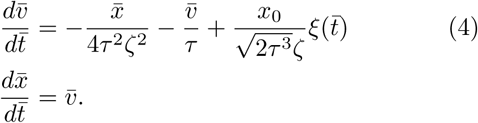

We make two changes of variable to simplify our expressions. We set 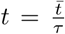 and 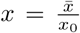. We further define a rescaled velocity, 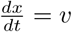, so that our equation of motion now reads

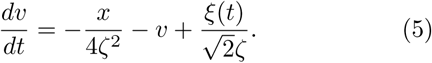

There are now two parameters that govern a particular solution to our information bottleneck problem: *ζ* and *∆t*, the timescale on which we want to retain optimal information about the future. We define *X*_*t*_ = (*x*(*t*), *v*(*t*)) and *X*_*t*+∆*t*_ = (*x*(*t* + ∆*t*), *v*(*t* + ∆*t*)) and seek a representation, 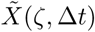, that can provide a maximum amount of information about *X*_*t*+∆*t*_ for a fixed amount of information about *X*_*t*_. We note that due to the Gaussian structure of the joint distribution of *X*_*t*_ and *X*_*t*+∆*t*_ for the SDDHO, the problem can be solved analytically. The optimal compressed representation is a noisy linear transform of *X*_*t*_ (see Appendix A) [25],

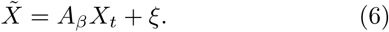

*A*_*β*_ is a matrix whose elements are a function of β, the tradeoff parameter in the information bottleneck objective function, and the statistics of the input and output variables. The added noise term, ξ, has the same dimensions as *X*_*t*_ and is a Gaussian variable with zero mean and unit variance.

We calculate the optimal compression, 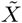, and its predictive information (see Appendix B.2). The past and future variables in the SDDHO bottleneck problem are jointly Gaussian, which means that the optimal compression can be summarized by its second-order statistics. We generalize analytically the results that were numerically obtained in Ref. [9] and explore the full parameter space of this dynamical model and examine all predictive bottleneck solutions, including different desired prediction timescales.

We quantify the efficiency of the representation 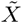 in terms of the variance of the following four probability distributions: the prior distribution, 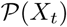, the distribution of the past conditioned on the compression, 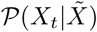, the distribution of the future conditioned the compressed variable 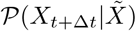, and the distribution of the future conditioned on exact knowledge of the past 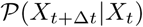. We represent the uncertainty reduction using two dimensional contour plots that depict the variances of the distributions in the ((*x* − 〈*x*〉)/*σ*_*x*_, (*v* − 〈*v*〉)/*σ*_*v*_) plane, where *σ*_*x*_ and *σ*_*v*_ are the standard deviations of the signal distribution 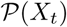.

The representation, 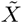, will be at most two-dimensional, with each of its components corresponding to linear combinations of position and velocity. It may be lower dimensional for certain values of *β*. The smallest critical *β* for which the representation remains two-dimensional is given in terms of the smallest eigenvalue *λ*_2_ of the matrix 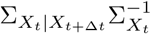 as *β*_*c*_ = 1/(1 − *λ*_2_) (see Appendix B.2). 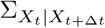 is the covariance matrix of the probability distribution of 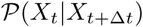 and 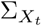 is the input variance. Below this critical β, the compressed representation is one dimensional, 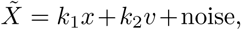, but it is still a combination of position and velocity.

Limiting cases along the the information bottleneck curve help build intuition about the optimal compression. If 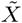 provides no information about the stimulus (e.g. *β* = 0), the variances of both of the conditional distributions match that of the prior distribution, 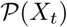, which is depicted as a circle of radius 1 (blue circle in Fig. 2b). However, if the encoding contains information about the past, the variance of 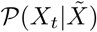 will be reduced compared to the prior. The maximal amount of predictive information, which is reached when *β* → ∞, can be visualized by examining the variance of 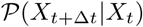 (e.g. the purple contour in Fig. 2b), which quantifies the correlations in *X*, itself, with no compression. Regardless of how precisely the current state of the stimulus is measured, the uncertainty about the future stimulus cannot be reduced below this minimal variance, because of the noise in the equation of motion.

From Figure 2b, we see that the conditional distribution 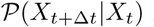 is strongly compressed in the position coordinate with some compression in the velocity coordinate. The information bottleneck solution at a fixed compression level (e.g. 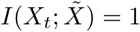) = 1), shown in Fig. 3a (left), gives an optimal encoding strategy for prediction (yellow curve) that reduces uncertainty in the position variable. This yields as much predictive information, 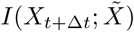, as possible for this value of 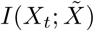. The uncertainty of the prediction is illustrated by the purple curve. We can explore the full range of compression levels, tracing out a full information bottleneck curve for this damping coefficient and desired prediction timescale, as shown in Figure 3. Velocity uncertainty is only reduced as we allow for less compression, as shown in Fig. 3a (right). For both of the cases represented in Fig. 3a, the illustrated encoding strategy yields a maximal amount of mutual information between the compressed representation, 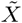, and the future for the given level of compression, as indicated by the red dots in Fig. 3b.

**FIG. 3:**
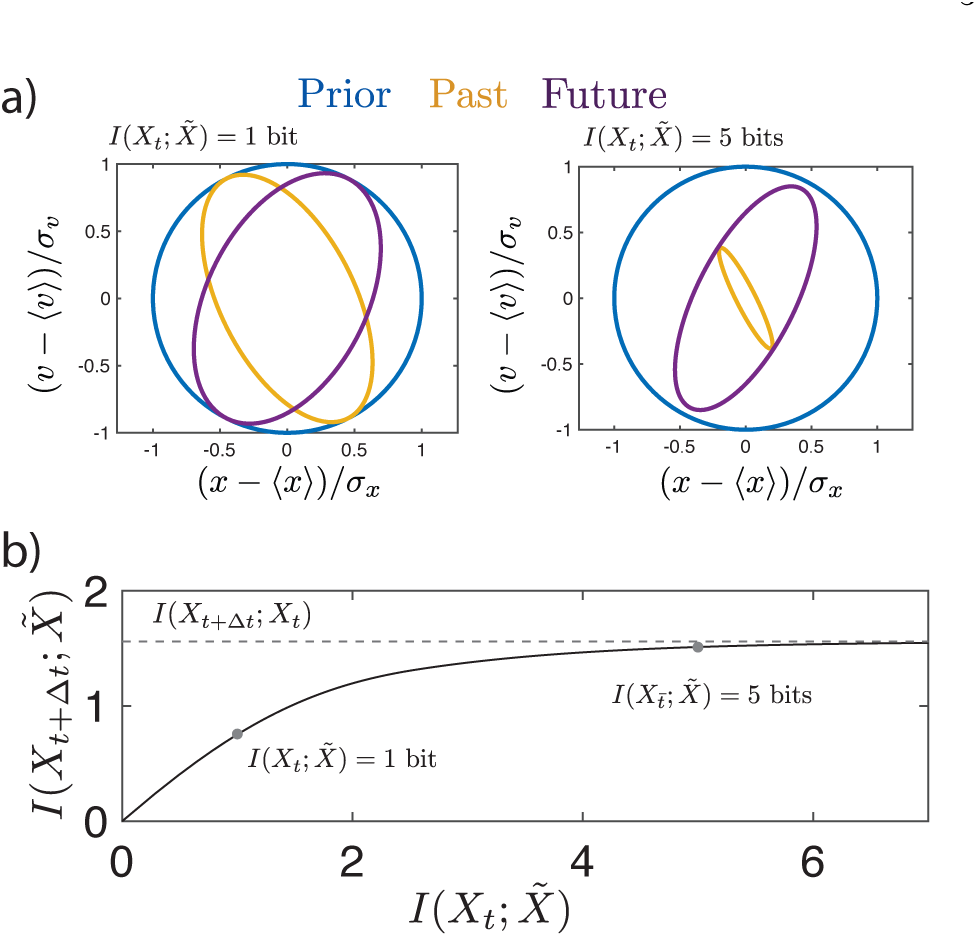
We consider the task of predicting the path of an SDDHO with 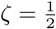 and ∆*t* = 1. (a) (left) We encode the history of the stimulus, *X*_*t*_, with a representation generated by the information bottleneck, 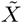, that can store 1 bit of information. Knowledge of the coordinates in the compressed representation space enables us reduce our uncertainty about the bar’s position and velocity, with a confidence interval given by ellipse in yellow. This particular choice of encoding scheme enables us to predict the future, *X*_*t*+∆*t*_ with a confidence interval given by the purple ellipse. The information bottleneck guarantees this uncertainty in future prediction is minimal for a given level of encoding. (right) The uncertainty in the prediction of the future can be reduced by reducing the overall level of uncertainty in the encoding of the history, as demonstrated by increasing the amount of information 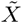 can store about *X*_*t*_. However, the uncertainty in the future prediction cannot be reduced below the variance of the propagator function. (b) We show how the information with the future scales with the information in the past, highlighting the points represented in panel (a).

As noted above, there is a phase transition along the information bottleneck curve, where the optimal, predictive compression of *X*_*t*_ changes from a one-dimensional representation to a two-dimensional one. This phase transition can be pinpointed in *β* for each choice of *ζ* and ∆*t*, and can be determined using the procedure described in is given in the Appendix A. To understand which directions are most important to represent at high levels of compression, we derive the analytic form of the leading eigenvector, *w*_1_, of the matrix 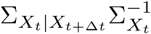. We have defined 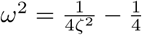 such that

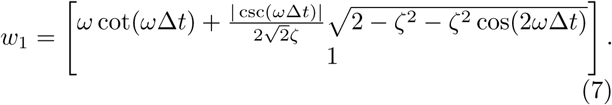

The angle of the encoding vector from the position direction is then given by

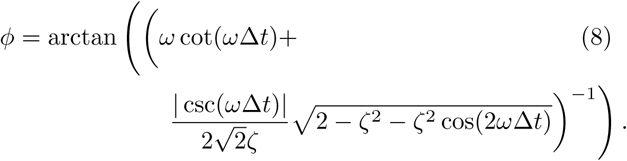

We consider *φ* in three limits: (I) the small ∆*t* limit, (II) the strongly overdamped limit (*ζ* → ∞), and (III) the strongly underdamped limit (*ζ* → 0).

(I): When *ω*∆*t* « 1, the angle can be expressed as

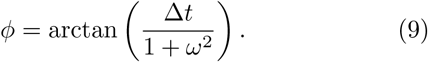

This suggests that for small *ω*∆*t*, the optimal encoding scheme favors position information over velocity information. The change in angle of the orientation from the position axis in this limit goes as *O*(∆*t*).

(II): The strongly overdamped limit. In this limit, *φ* becomes

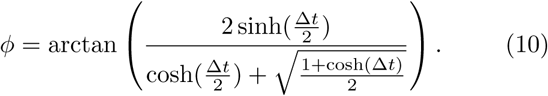

In the large ∆*t* limit, 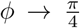. In the small ∆*t* limit, *φ* → arctan(∆*t*). Past position information is the best predictor of the future input at short lags, which velocity and position require equally fine representation for prediction at longer lags.

(III) The strongly underdamped limit. In this limit, *φ* can be written as

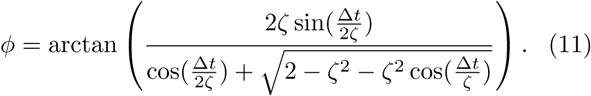

We observe periodicity in the optimal encoding angle between position and velocity. This means that the optimal tradeoff between representing position or velocity depends on the timescale of prediction. However, the denominator never approaches 0, so the encoding scheme never favors pure velocity encoding. It returns to position-only encoding when ∆*t*/2*ζ* = *nπ*.

At large compression values, i.e. small amounts of information about the past, the information bottleneck curve is approximately linear. The slope of the information bottleneck curve at small 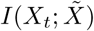 is given by 1 − *λ*_1_, where λ_1_ is the smallest eigenvalue of the matrix, 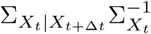. The value of the slope is

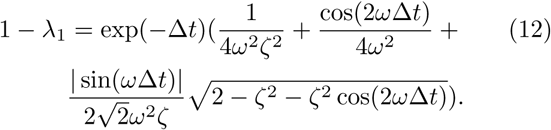

For large ∆*t*, it is clear that the slope will be constrained by the exponential term, and the information will fall as exp(−∆*t*) as we attempt to predict farther into the future. For small ∆*t*, however, we see that the slope goes as 1 − ∆*t*^2^, and our predictive information decays more slowly.

For vanishingly small compression, i.e. *β* → ∞, the predictive information that can be extracted by 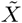 approaches the limit set by the temporal correlations in *X*, itself, given by

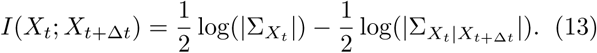

For large ∆*t*, this expression becomes

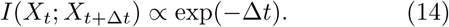

For small ∆*t*,

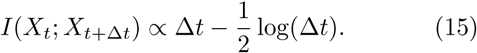

The constants emerge from the physical parameters of the input dynamics.

#### 1. Optimal representations in all parameter regimes for fixed past information

We sweep over all possible parameter regimes of the SDDHO keeping 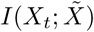 fixed to 5 bits and find the optimal representation for a variety of timescales (Fig. 4), keeping a fixed amount of information encoded about the past for each realization of the stimulus and prediction. More information can be transmitted for shorter delays (Fig. 4a,d,g) between the past and future signal than for longer delays (Fig. 4c,f,i). In addition, at shorter prediction timescales more information about the past is needed to reach the upper bound, as more information can be gleaned about the future. In particular, for an overdamped SDDHO at short timescales (Fig. 4a), the evolution of the equations of motion are well approximated by integrating Eq. 3 with the left hand side set to zero, and the optimal representation encodes mostly positional information. This can be observed by noting that the encoding ellipse remains on-axis and mostly compressed in the position dimension. For the underdamped case, in short time predictions (Fig. 4g), a similar strategy is effective. However, for longer predictions (Fig. 4h,i), inertial effects cause position at one time to be strongly predictive of future velocity and vice versa. As a result, the encoding distribution has to take advantage of these correlations to be optimally predictive. These effects can be observed in the rotation of the encoding ellipse, as it indicates that the uncertainty in position-velocity correlated directions are being reduced, at some cost to position and velocity encoding. The critically damped SDDHO (Fig. 4d-f) demonstrates rapid loss of information about the future, like that observed in the underdamped case. The critically damped case displays a bias towards encoding position over velocity information at both long and intermediate timescales, as in the overdamped case. At long timescales, Fig. 4f, the optimal encoding is non-predictive.

**FIG. 4:**
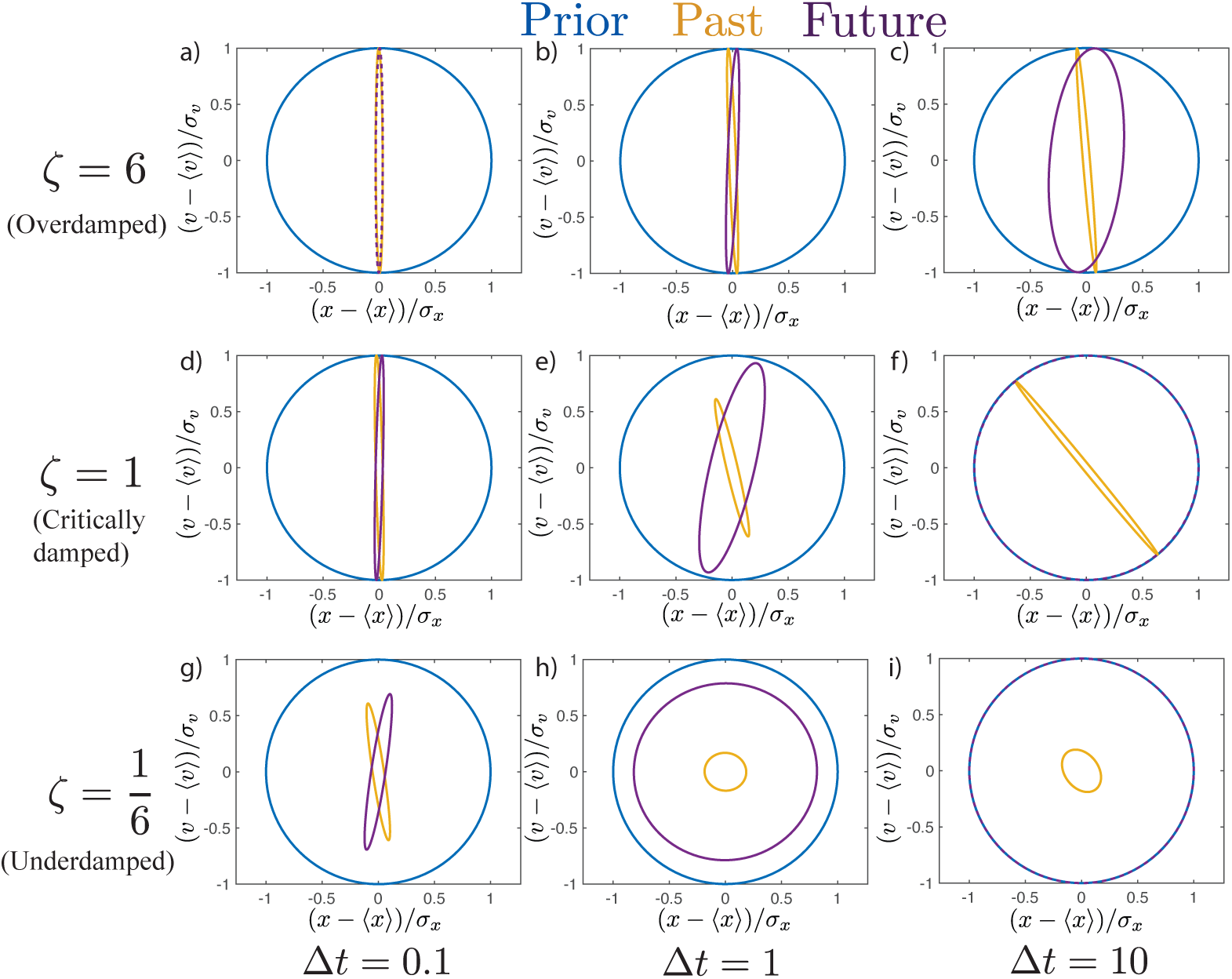
Possible behaviors associated for the SDDHO for a variety of timescales with a fixed 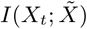 of 5 bits. For an overdamped SDDHO, panel a-c, the optimal representation continues to encode mostly position information, as velocity is hard to predict. For the underdamped case, panels g-i, as the timescale of prediction increases, the optimal representation changes from being mostly position information to being a mix of position and velocity information. Optimal representations for critically damped input motion are shown in panels d-f. Comparatively, overdamped stimuli do not require precise velocity measurements, even at long timescales. Optimal predictive representations of overdamped input dynamics have higher amounts of predictive information for longer timescales, when compared to underdamped and critically damped cases.

#### 2. Suboptimal representations

Biological systems might not adapt to each input regime perfectly, nor may they be optimally efficient for every possible kind of input dynamics. We consider what happens when an optimal representation is changed, necessarily making it suboptimal for predicting the future stimulus. We construct a new representation by rotating the optimal solution in the position, velocity plane. We examine the conditional distributions for this suboptimal representation, both about the past, 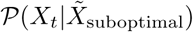, and the future, 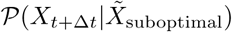. For a fixed amount of information about the past, 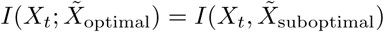, we compare the predictive information in the optimal (Fig. 5a) and the sub-optimal representations (Fig. 5b). In this example, we are exploring the impact of encoding velocity with high certainty as compared to encoding position with high certainty. We observe that encoding velocity provides very little predictive power, indicating that encoding velocity and position is not equally important, even for equal compression levels. In addition, it shows that encoding schemes discovered by IB are optimal for predictive purposes.

**FIG. 5:**
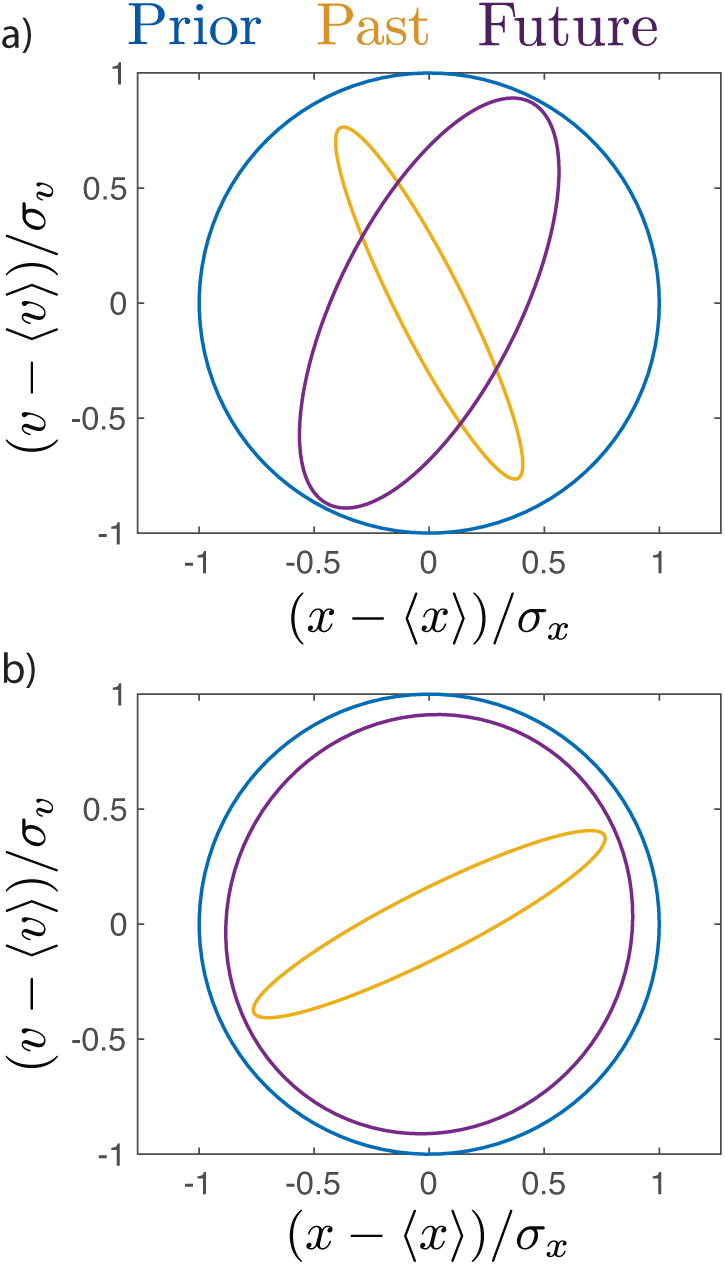
Example of a sub-optimal compression. An optimal predictive, compressed representation, in panel (a) compared to a suboptimal representation, in panel (b) for a prediction of ∆*t* = 1 away in the underdamped regime (*ζ* = 1/2). We fix the mutual information between the representations and the past 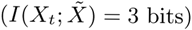, but find that, as expected, the sub-optimal representation contains significantly less information about the future.

#### 3. Kalman filters versus information bottleneck

We can also compare our information bottleneck solutions to what one would obtain using Kalman filters [26]. We note that Kalman filters are not designed to be *efficient* strategies for extracting predictive information, as shown in the Appendix, Figure B.1. This is because the Kalman filter approach does not constrain the representation entropy (i.e. it does not have a resource-limit constraint). A Kalman filter also always explicitly makes a model of the dynamics that generate updates to the input variable, an explicit model of the ‘physics of the external world’. The information bottleneck framework enables exploration of representations without explicitly developing an internal model of the dynamics and also includes resource constraints. Thus, for a given amount of compression, the information bottleneck solution to the prediction problem is as predictive as possible, whereas a Kalman filter may miss important predictive features of the input while representing noisy, unpredictable features. In that sense, the Kalman filter approach is agnostic about what input bits matter for prediction, and is a less efficient coding scheme of predictive information for a given channel capacity.

#### 4. Transferability of a representation

So far, we have described the form that optimal predictive compressions take along the information bottleneck curve for a given *ζ* and ∆*t*. How do these representations translate when applied to other prediction timescales (i.e. can the optimal predictive scheme for near-term predictions help generate long-term predictions, too?) or other parameter regimes of the model? This may be important if the underlying parameters in the external stimulus are changing rapidly in comparison to the adaptation timescales in the encoder, which we imagine to be a biological network. One possible solution is for the encoder to employ a representation that is useful across a wide range of input statistics. This requires that the predictive power of a given representation is, to some extent, transferrable to other input regimes. To quantify how ‘transferrable’ different representations are, we take an optimal representation from one (ζ, ∆*t*) and ask how efficiently it captures predictive information for a different parameter regime, (*ζ′*, ∆*t′*).

We identify these global strategies by finding the optimal encoder for a stimulus with parameters (*ζ*, ∆*t*) that generates a representation, 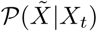, at some given compression level, *I*_past_. We will label the predictive information captured by this representation 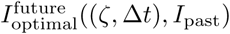. We hold the representation fixed and apply it to a stimulus with different underlying parameters (*ζ′*, ∆*t′*) and compute the amount of predictive information the previous representation yields for this stimulus. We call this the transferred predictive information 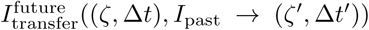. We note that 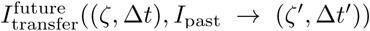, *I*_past_ → (*ζ′*, ∆*t′*)) may sometimes be larger than 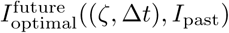, because changing (ζ, ∆*t*) may increase both *I*_past_ and *I*_future_ (see e.g. Figure 6a).

**FIG. 6:**
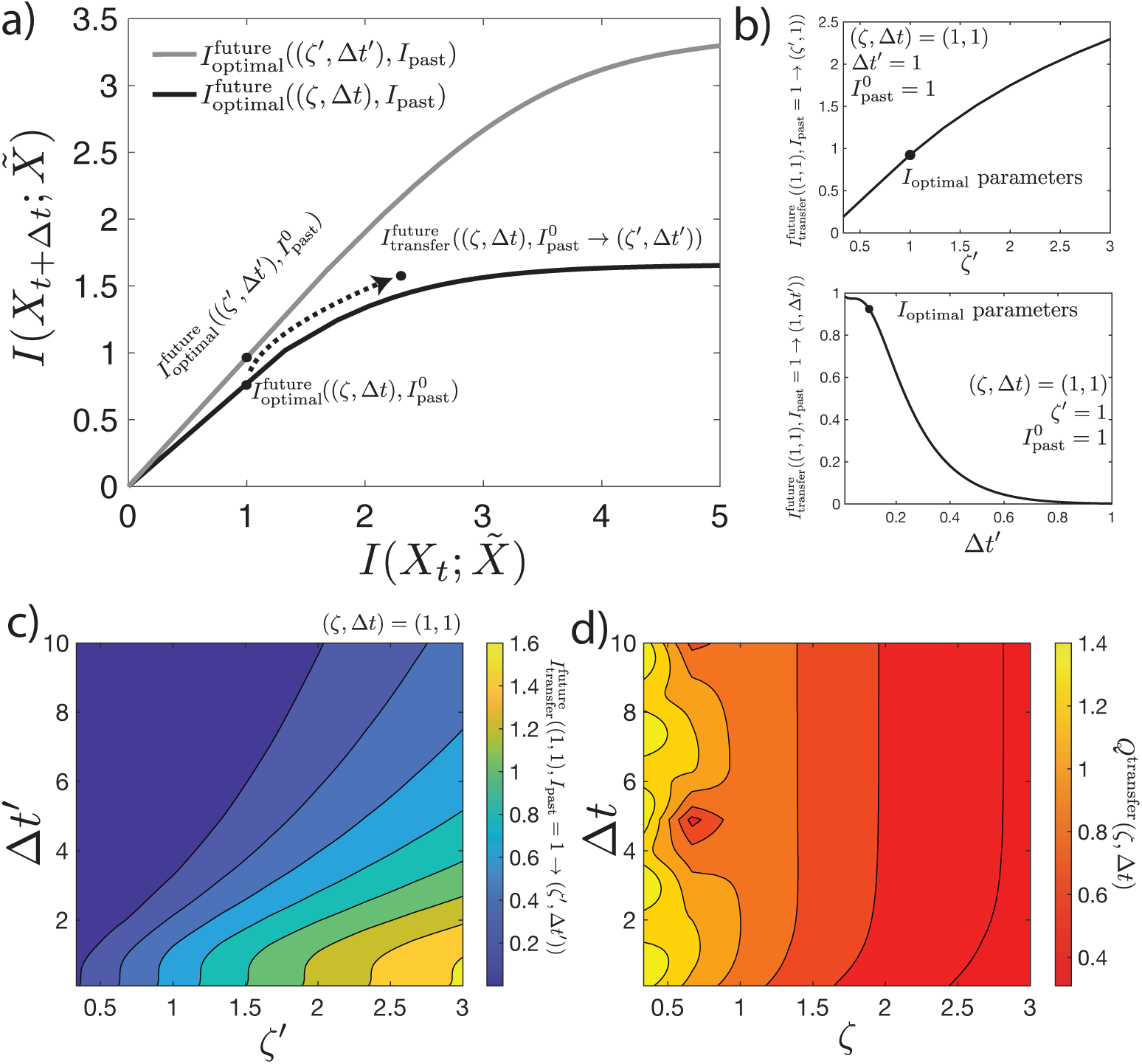
Representations learned on underdamped systems can be transferred to other types of motion, while representations learned on overdamped systems cannot be easily transferred. (a) Here, we consider the information bottleneck bound curve (black) for a stimulus with underlying parameters, (*ζ*, ∆*t*). For some particular level of 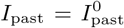, we obtain a mapping, 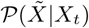 that extracts some predictive information, denoted 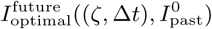, about a stimulus with parameters (*ζ*, ∆*t*). Keeping that mapping fixed, we determine the amount of predictive information for dynamics with new parameters (*ζ′*, ∆*t′*), denoted by 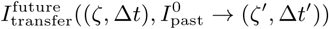. (b) One-dimensional slices of 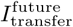 in the (*ζ′*, ∆*t′*) plane: 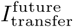 versus *ζ′* for ∆*t′* = 1. 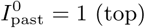, and versus ∆*t′* for *ζ′* = 1. Parameters are set to (*ζ* = 1, ∆*t* = 1), 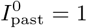. (c) Two-dimensional map of 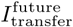 versus (*ζ′*, ∆*t′*) (same parameters as b). (d) Overall transferability of the mapping. The heatmap of (c) is integrated over *ζ′* and ∆*t′* and normalized by the integral of 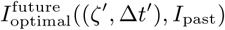. We see that mappings learned from underdamped systems at late times yield high levels of predictive information for a wide range of parameters, while mappings learned from overdamped systems are not generally useful.

For every fixed (ζ, ∆*t*) and *I*_past_, we can take the optimal 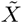 and transfer it to a wide range of new *ζ′*’s and timescales, ∆*t′*. For a particular example (*ζ*, ∆*t*), this is shown in Figure 6b. The representation optimized for critical damping is finer-grained than what’s required in the overdamped regime. We can sweep over all combinations of the new *ζ′*,’s and ∆*t′*s. What we get, then, is a mapping of 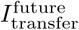 for this representation that was optimized for one particular (*ζ*, ∆*t*) pair across all new (*ζ′*, ∆*t′*)’s. This is shown in Figure 6c, (Figure 6b are just two slices through this surface). This surface gives a qualitative picture the transferability of this particular representation.

To get a quantitative summary of this behavior that we can then compare across different starting points (*ζ*, ∆*t*), we integrate this surface over 1/3 < *ζ′* < 3, 0.1 < ∆*t′* < 10, and then normalize by the integral of 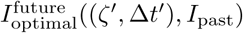 over the same surface. This yields an overall transferability measure, *Q*^transfer^(*ζ*, ∆*t*). We report these results in Figure 6d. Representations that are optimal for underdamped systems at late times are the most transferable. This is because generating a predictive mapping for underdamped motion requires some measurement of velocity, which is generally useful for many late-time predictions. Additionally, prediction of underdamped motion requires high precision measurement of position, and that information is broadly useful across all parameters.

### B. History-dependent Gaussian Stimuli

In the above analysis, we considered stimuli with correlations that fall off exponentially. However, natural scenes, such as leaves blowing in the wind or bees moving in their hives, are shown to have heavy-tailed statistics [21, 27, 28], and we extend our results to models of motion stimuli with heavy-tailed temporal correlation. Despite long-ranged temporal order, prediction is still possible. We show this through the use of the Generalized Langevin equation [29–31]:

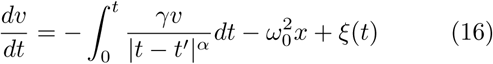

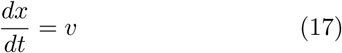

Here, we have returned to unscaled definitions of *v*, and *t*. The damping force here is a power-law kernel. In order for the system to obey the fluctuation-dissipation theorem, we note that *〈ξ*(*t*)〉 = 0, and 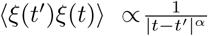. In this dynamical system, position autocorrelation *〈x*(*t*)*x*(*t*′)〉 ∼ *t*^*α*^ and velocity autocorrelation *〈v*(*t*)*v*(*t′*)〉 ∼ *t*^−*α*−1^ for large *t*.

The prediction problem is similar to the prediction problem for the memoryless SDDHO, but we now take an extended past, 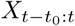, for prediction of an extended future, 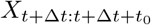, where *t*_0_ sets the size of the window into the past we consider and the future we predict (Fig. 7a). Using the approach described in Appendix A, we compute the optimal representation and determine how informative the past is about the future. The obajective function for this extended information bottleneck problem is,

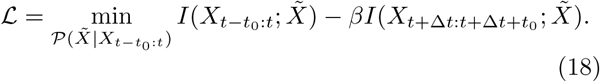

**FIG. 7:**
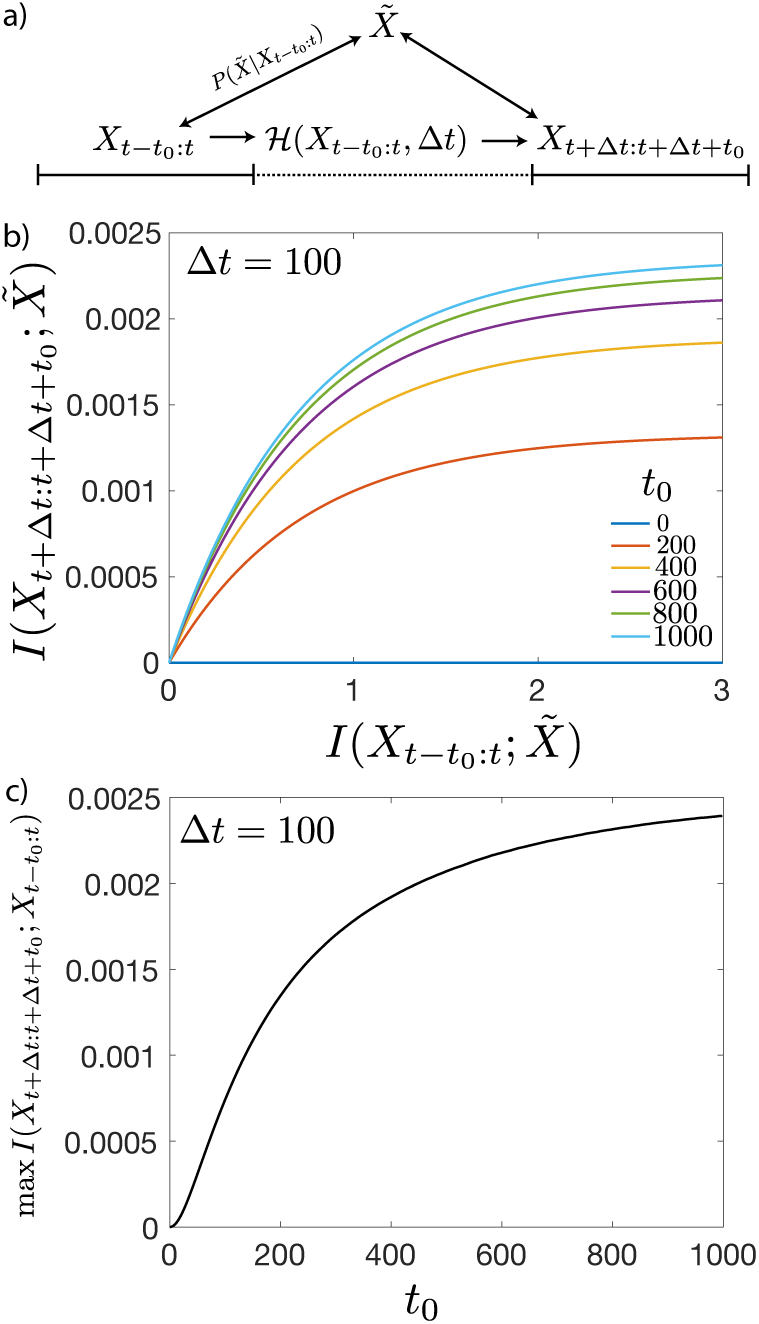
The ability of the information bottleneck Method to predict history-dependent stimuli. (a) The prediction problem, using an extended history and a future. This problem is largely similar to the one set up for the SDDHO but the past and the future are larger composites of observations within a window of time *t* − *t*_0_: *t* for the past and *t*+∆*t*: *t*+∆*t*+*t*_0_ for the future. (b) Predictive information 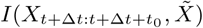 with lag ∆*t*. (c) The maximum available predictive information saturates as a function of the historical information used *t*_0_.

The information bottleneck curves show more predictive information as the prediction process uses more past information (larger *t*_0_ in Fig. 7b). Not including any history results in an inability to extract the predictive information. However, for low compression, large *β*, we find that the amount of predictive information that can be extracted saturates quickly as we increase the amount of history, *t*_0_. This implies diminishing returns in prediction for encoding history. Despite the diverging autocorrelation timescale, prediction only functions on a limited timescale and the maximum available prediction information always saturates as a function of *t*_0_ (Fig. 7c). These results indicate that efficient coding strategies can enable prediction even in complex temporally correlated environments.

### C. Evolutionary dynamics

Exploiting temporal correlations to make predictions is not limited to vision. Another aspect of the prediction problem appears in the adaptive immune system, where temporal correlations in pathogen evolution may be exploited to help an organism build up an immunity. Exploiting these correlations can be done at a population level, in terms of vaccine design [32–35], and has been postulated as a means for the immune system to adapt to future threats [11, 36]. Here, we present efficient predictive coding strategies for the Wright-Fisher model, which is commonly used to describe viral evolution [37]. In contrast to the two models studied so far, Wright-Fisher dynamics are not Gaussian. We use this model to explore how the results obtained in the previous sections generalize to non-Gaussian statistics of the past and future distributions.

Wright-Fisher models of evolution assume a constant population size of *N*. We consider a single mutating site with each individual in the population having either a wild-type or a mutant allele at this site. The allele choice of subsequent generations depends on the frequency of the mutant allele in the ancestral generation at time t, *X*_*t*_, the selection pressure on the mutant allele, *s*, and the mutation rate from the wild-type to the mutant allele and back, *µ*, as depicted as Fig. 8a. For large enough *N*, the update rule of the allele frequencies is given through the diffusion approximation interpreted with the Ito convention [38]:

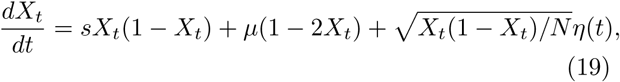

where *〈η*(*t*)〉 = 0, *〈η*(*t*)*η*(*t′*)〉 = *δ*(*t* − *t′*).

**FIG. 8:**
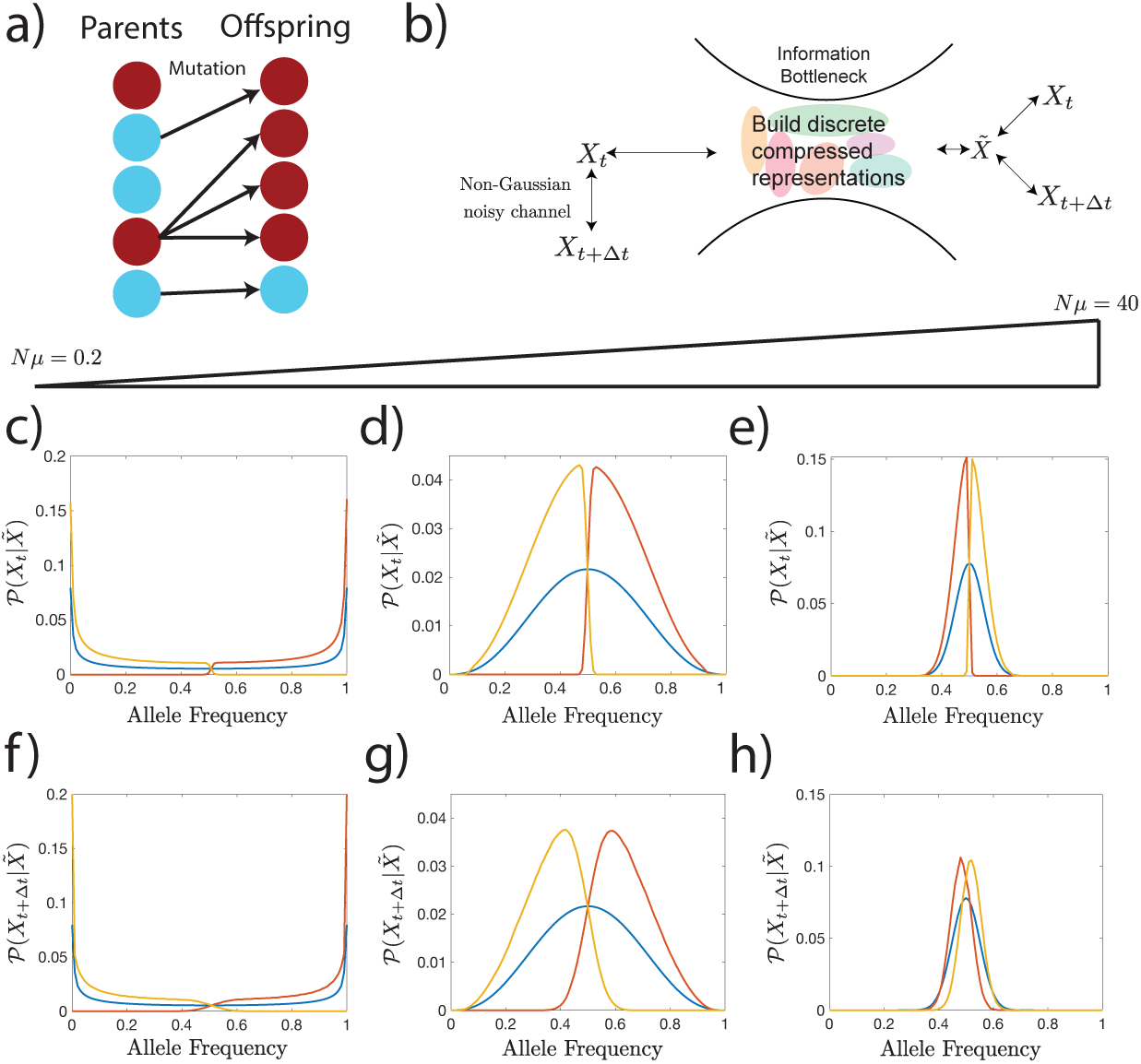
The information bottleneck solution for a Wright Fisher process. (a) The Wright-Fisher model of evolution can be visualized as a population of *N* parents giving rise to a population of *N* children. Genotypes of the children are selected as a function of the parents’ generation genotypes subject to mutation rates, *µ*, and selective pressures *s*. (b) Information bottleneck schematic with a discrete (rather than continuous) representation variable, 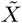. (c-h) We explore information bottleneck solutions to Wright-Fisher dynamics under the condition that the cardinality of 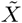, *m*, is 2 and take *β* to be large enough that 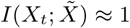, *β* ≈ 4. Parameters: *N* = 100, *N*_*s*_ = 0.001, ∆*t* = 1, and *N*_*µ*_ = 0.2, *N*_*µ*_ = 2, and *N*_*µ*_ = 40 (from left to right). (c-e) In blue, we plot the steady state distribution. In yellow and red, we show the inferred historical distribution of alleles based on the observed value of 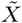. Note that each distribution is corresponds to roughly non-overlapping portions of allele frequency space. (f-h) Predicted distribution of alleles based on the value of 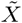. We observe that as mutation rate increases, the timescale of relaxation to steady state decreases, so historical information is less useful and the predictions becomes more degenerate with the steady state distribution.

For this model, defining the representation 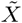 as a noisy linear transformation of the past frequency *X*_*t*_ as we did for the Gaussian case in Eq. 21 does not capture all of the dependencies of the future on the past due to the non-Gaussian character of the joint distribution of *X*_*t*+∆*t*_ and *X*_*t*_ stemming from the non-linear form of Eq. 19. Instead, we determine the mapping of *X*_*t*_ to 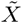 numerically using the Blahut-Arimoto algorithm [39, 40]. For ease of computation, we will take the representation variable 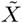 to be discrete (Fig. 8b) and later, approximate continuous 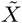 by driving the cardinality of 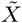, denoted by *m*, to be high. The assumption that 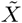 is discrete results in each realization of the representation tiling a distinct part of frequency space. This encoding scheme can be thought of as lymphocyte antigen-receptors in the adaptive immune system corresponding to different regions of phenotypic space [41].

We first consider the example with *m* = 2 representations. In the weak mutation, weak selection limit (*N*_*µ*_, *N*_*s*_ ≪ 1), the steady state probability distribution of allele frequencies,

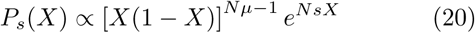

(blue line in Fig. 8c) is peaked around the frequency boundaries, indicating that at long times, an allele either fixes or goes extinct. In this case, one value of the representation variable corresponds to the range of high allele frequencies and the other corresponds to low allele frequencies (Fig. 8c, yellow and red lines). These encoding schemes can be used to make predictions, whether it be by an observer or the immune system, via determining the future probability distribution of the alleles conditioned on the value of the representation variables, 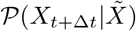. We present these predictions in Fig. 8f. The predictive information conferred by the representation variable is limited by the information it has about the past, as in the Gaussian case (Fig. 10a.)

For larger mutation rates, the steady state distribution becomes centered around the equal probability of observing either one of the two alleles, but the two representation variables still cover the frequency domain in way that minimizes overlap (Fig. 8d and e). We observe a sharp drop in 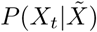 at the boundary between the two representations. The future distribution of allele frequencies in this region (Fig. 8g and h), however, displays large overlap. The degree of this overlap increases as the mutation rate gets larger, suggesting prediction is harder in the strong mutation limit. The optimal encoding of the past distribution biases the representation variable towards frequency space regions with larger steady state probability mass.

In Fig. 9, we explore the consequence of transferring a mapping, 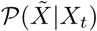), from a high mutation model to a low mutation model and vice versa. We observe that the weak mutation representation is more transferrable than the strong mutation representation. One reason for this is that the strong mutation limit provides little predictive information, as seen in Fig. 10b. In addition, high mutation representations focus on *X* = 1/2, while the population more frequently occupies allele frequencies near 0 and 1 in other regimes. Comparatively, representations learned on weak mutation models can provide predictive information, because they cover more evenly the spectrum of allele frequencies.

**FIG. 9:**
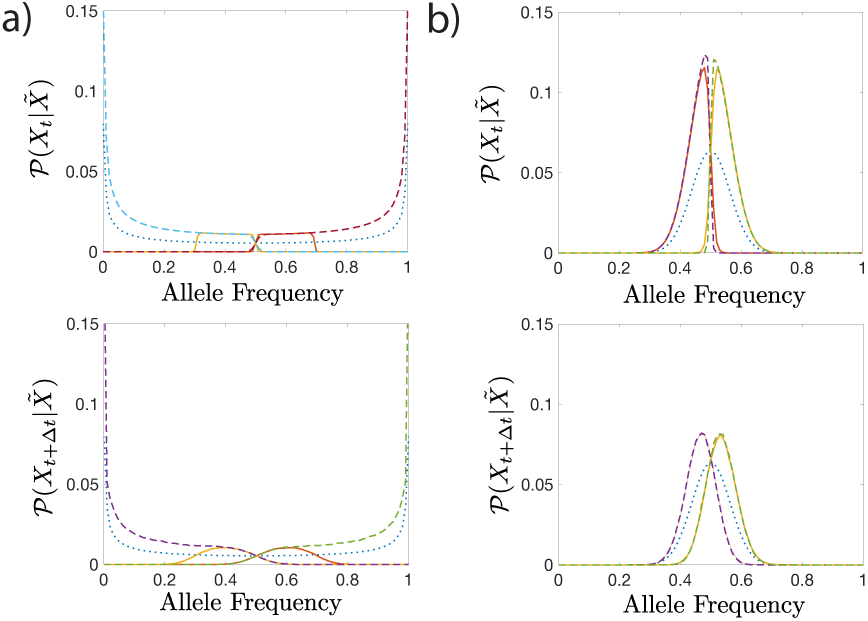
Transferability of prediction schemes in Wright-Fisher dynamics. We transfer a mapping, 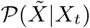, trained on one set of parameters and apply it to another. We consider transfers between two choices of mutability, 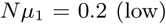 and 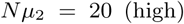, with *N* = 100, *N*_*s*_ = 0.001, ∆*t* = 1. The dotted line is the steady state allele frequency distribution, the solid lines are the transferred representations, and the dashed lines are the optimal solutions. The top panels correspond to the distributions of *X*_*t*_ and the bottom panels correspond to distributions of *X*_*t*+∆*t*_. (a) Transfer from high to low mutability. Optimal information values: 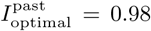 and 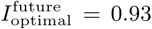; transferred information values: 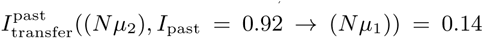 and 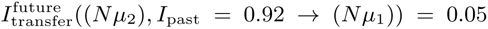. Representations learned on high mutation rates are not predictive in the low mutation regime. (b) Transfer from low to high mutability. Optimal information values: 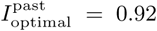 and 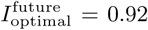 and 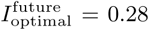. Transferred information values: 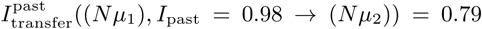 and 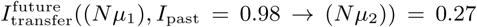. Transfer in this direction yields good predictive informations.

**FIG. 10:**
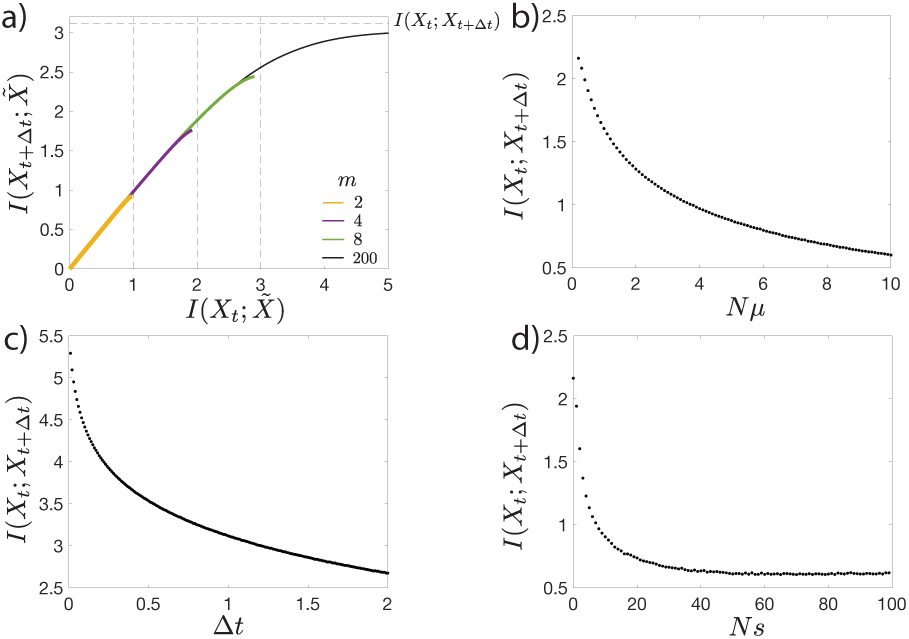
Amount of predictive information in the Wright Fisher dynamics as a function of the model parameters. (a) Predictive information as a function of compression level. Predictive information increases with the cardinality, *m*, of the representation variable. The amount of predictive information is limited by log(*m*) (vertical dashed lines) for small *m*, and the mutual information between the future and the past, *I*(*X*_*t*+∆*t*_; *X*_*t*_) (horizontal dashed line), for large *m*. Bifurcations occur in the amount of predictive information. For small 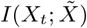, the encoding strategies for different *m* are degenerate and the degeneracy is lifted as 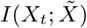 increases, with large *m* schemes accessing higher 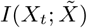 ranges. Parameters: *N* = 100, *N*_*µ*_ = 0.2, *N*_*µ*_ = 0.2, *N*_*s*_ = 0.001, ∆*t* = 1. (b-d), Value of the asymptote of the information bottleneck curve, *I*(*X*_*t*_; *X*_*t*+∆*t*_) with: (b) *N* = 100, *N*_*s*_ = 0.001, ∆*t* = 1 as a function of *µ*; (c) *N* = 100, *N*_*µ*_ = 0.2, *N*_*s*_ = 0.001 as a function of ∆*t*; and (d) *N* = 100, *N*_*µ*_ = 0.2, and ∆*t* = 1 as a function of *s*.

We can extend the observations in Fig. 8 to see how the predictive information depends on the strength of the selection and mutation rates (Fig. 10b and d). Prediction is easiest in the weak mutation and selection limit, as population genotype change occur slowly and the steady state distribution is localized in one regime of the frequency domain. For evolutionary forces acting on faster timescales, prediction becomes harder since the relaxation to the steady state is fast. Although the mutation result might be expected, the loss of predictive information in the high selection regime seems counterintuitive: due to a large bias between one of the two alleles evolution appears reproducible and “predictable” in the high selection limit. This bias renders the allele state easier to guess but this is not due to information about the initial state. The mutual information-based measure of predictive information used here captures a reduction of entropy in the estimation of the future distribution of allele frequencies due to conditioning on the representation variable. When the entropy of the future distribution of alleles *H*(*X*_*t*+∆*t*_) is small, the reduction is small and predictive information is also small. As expected, predictive information decreases with time ∆*t*, since the state *X*_*t*_ and *X*_*t*+∆*t*_ decorrelate due to noise (Fig. 10c).

So far we have discussed the results for *m* = 2 representations. As we increase the tradeoff parameter, *β* in Eq. 1, the amount of predictive information increases, since we retain more information about the past. However, at high *β* values the amount of information the representation variable can hold saturates, and the predictive information reaches a maximum value (1 bit for the *m* = 2 yellow line in Fig. 10a). Increasing the number of representations *m* to 3 increases the range of accessible information the representation variable has about the past *I*(*X*_*t*_; *X*), increasing the range of predictive information (purple line in Fig. 10a)). Comparing the *m* = 2 and *m* = 3 representations for maximum values of *β* for each of them (Fig. 11a and b), shows that larger numbers of representations tile allele frequency space more finely, allowing for more precise encodings of the past and future distributions. The maximum amount of information about the past goes as log(*m*) (Fig. 10a). The predictive information curves for different *m* values are the same, until the branching point ≲ log(*m*) for each *m* (Fig. 10a).

**FIG. 11:**
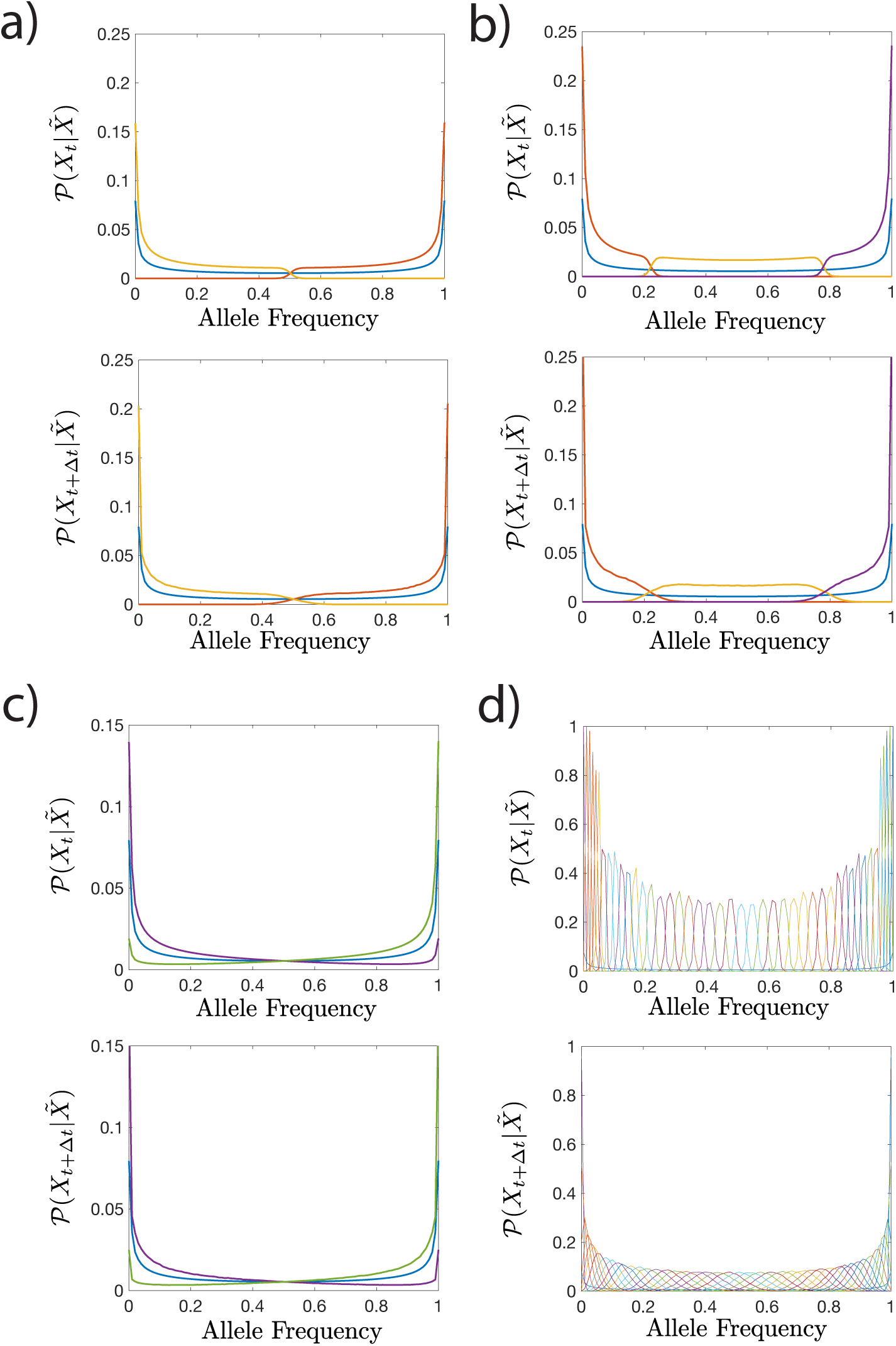
Encoding schemes with *m* > 2 representation variables. The representations which carry maximum predictive information for *m* = 2 at 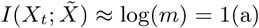 and *m* = 3 at 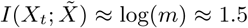. (b). The optimal representations at large *m* tile space more finely and have higher predictive information. The optimal representations for *m* = 200 at fixed *β* = 1.01 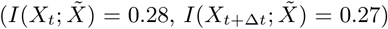 (c) and *β* = 20 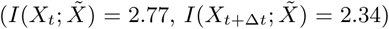. (d) At low 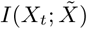), many of the representations are redundant and do not confer more predictive information than the *m* = 2 scheme. A more explicit comparison is given in Appendix Fig. C.2. At high 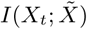, the degeneracy is lifted. All computations done at *N* = 100, *N*_*µ*_ = 0.2, *N*_*s*_ = 0.001, ∆*t* = 1.

We analyze the nature of this branching by taking *m* ≫ 1, *m* = 200 (Fig. 11c and d). At small *β* (and corresponding small 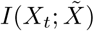) the optimal encoding scheme is the same if we had imposed a small *m* (Fig. 11c), with additional degenerate representations (Fig. C.2). By increasing *β* (and 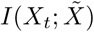), the degeneracy is lifted and additional representation cover non-overlapping regimes of allele frequency space. This demonstrates the existence of a critical *β* for each predictive coding scheme, above which *m* needs to be increased to extract more predictive information and below which additional values of the representation variable encode redundant portions of allele frequency space. While we do not estimate the critical *β*, approaches to estimating them are presented in [42, 43].

The *m* = 200 encoding approximates the continuous 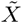 representation. In the high 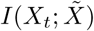 limit, the *m* = 200 encoding gives precise representations (i.e. with low variability in 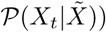 in regions of allele frequency space with high steady state distribution values, and less precise representations elsewhere (Fig. 11d top panel, Fig. C.3). This dependence differs from the Gaussian case, where the uncertainty of the representation is independent of the encoded value. The decoding distributions 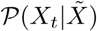 are also not Gaussian. This encoding builds a mapping of internal response to external stimuli, by tiling the internal representation space of external stimuli in a non-uniform manner. These non-uniform frequency tilings are similar to Laughlin’s predictions for maximally informative coding in vision [2], but with the added constraint of choosing the tiling to enable the most informative predictions.

## III. DISCUSSION

We have demonstrated that the information bottleneck method can be used to construct predictive encoding schemes for a variety of biologically-relevant dynamic stimuli. The approach described in this paper can be used to make predictions about the underlying encoding schemes used by biological systems that are compelled by their behavioral and fitness constraints to make predictions. These results thus provide experimentally testable hypotheses. The key principle is that not all input dimensions are equally relevant for prediction; information encoding systems must be able to parse which dimensions are relevant when coding capacity is small relative to the available predictive information. Hence, the biological (or engineered) system must navigate a tradeoff between reducing the overall uncertainty in its prediction while only being able to make measurements with some fixed uncertainty.

We hypothesize that biological systems that need to operate flexibly across a wide range of different input statistics may use a best-compromise predictive encoding of their inputs. We have used a transferability metric, *Q*, to quantify just how useful a particular scheme is across other dynamic regimes and prediction timescales. What we have shown is that a compromise between representing position and velocity of a single object provides a good, general, predictor for a large set of input behaviors. When adaptation is slower than the timescale over which the environment changes, such a compromise might be beneficial to the biological system. On the other hand, if the biological encoder can adapt, the optimal predictive encoder for those particular dynamics is the best encoder. We have provided a fully-worked set of examples of what those optimal encoders look like for a variety of parameter choices. The dynamics of natural inputs to biological systems could be mapped onto particular points in these dynamics, providing a hypothesis for what optimal prediction would look like in that system.

We also explored the ability to predict more complex, non-Markovian dynamics. We asked about the usefulness of storing information about the past in the presence of power-law temporal correlations. The optimal information bottleneck solution showed fast diminishing returns as it was allowed to dig deeper and deeper into the past, suggesting that simple encoding schemes with limited temporal span have good predictive power even in complex correlated environments.

Superficially, our framework may seem similar to a Kalman filter [26]. There are few major differences in this approach. Kalman filtering algorithms have been used to explain responses to changes in external stimuli in biological system [44]. In this framework, the Kalman filters seek to maximize information by minimizing the variance of the true coordinates of an external input and the estimate of those coordinates. The estimate is, then, a prediction of the next time step, and is iteratively updated. Our information bottleneck approach extracts past information, but explicitly includes another constraint: resource limitations. The tuning of *I*_past_ is the main difference between our approach and a Kalman filter. Another major difference is that we do not assume the underlying encoder has any explicit representation of the ‘physics’ of the input. There is no internal model of the input stimulus, apart from our probabilistic mapping from the input to our compressed representation of that input. A biological system could have such an internal model, but that would add significant coding costs that would have to be treated by another term in our framework to draw a precise equivalence between the approaches. We show in the Appendix that the Kalman filter approach is not as efficient, in general, as the predictive information bottleneck approach that we present here.

The evolutionary context shows another set of solutions to predictive information in terms of discrete representations that tile input space. Although we impose discrete representations, their non-overlapping character remains even it the limit of many representations. These kinds of solutions are reminiscent of the Laughlin solution for information maximization of input and output in the visual system given a nonlinear noisy channel [2], since input space is covered proportionally to the steady state distribution at a given frequency. Tiling solutions have also been described when optimizing information in gene regulatory networks with nonlinear input-output relations, when one input regulates many gene outputs [45]. In this case each gene was expressed in a different region of the input concentration domain. Similarly to our example, where the lifting the degeneracy between multiple representations covering the same frequency range allows for the prediction of more information about the future, lifting the degeneracy between different genes making the same readout, increases the transmitted information between the input concentration and the outputs. More generally, discrete tiling solutions are omnipresent in information optimization problems with boundaries [46, 47].

Biologically, predicting evolutionary dynamics is a different problem than predicting motion. Maybe the accuracy of prediction matters less, while covering the space of potentially very different inputs is important. In our simple example, this is best seen in the strong mutation limit where the mutant allele either fixes or goes extinct with equal probability. In this case, a single Gaussian representation cannot give a large values of predictive information. A discrete representation, which specializes to different regions of input space, is a way to maximize predictive power for very different inputs. It is likely that these kinds of solutions generalize to the case of continuous, multi-dimensional phenotypic spaces, where discrete representations provides a way for the immune system to hedge its bets against pathogens by covering the space of antigen recognition[24]. The tiling solution that appears in the non-Gaussian solution of the problem is also potentially interesting for olfactory systems. The number of odorant molecules is much larger than odor receptors [48, 49], which can be thought of as representation variables that cover the phenotypic input space of odorants. The predictive information bottleneck solution gives us a recipe for covering space, given a dynamical model of evolution of the inputs.

The results in the non-Gaussian problem are different than the Gaussian problem in two important ways: the encoding distributions are not Gaussian (e.g. Fig. 8d and e), and the variance of the encoding distributions depends on the the value of 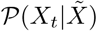 (Fig. 11d). These solutions offer more flexibility for internal encoding of external signals.

The information bottleneck approach has received a lot of attention in the machine learning community lately, because it provides a useful framework for creating well-calibrated networks that solve classification problems at human-level performance[14, 50, 51]. In these deep networks, variational methods approximate the information quantities in the bottleneck, and have proven their practical utility in many machine learning contexts. These approaches do not always provide intuition about how the networks achieve this performance and what the IB approach creates in the hidden encoding layers. Here, we have worked through a set of analytically tractable examples, laying the groundwork for building intuition about the structure of IB solutions and their generalizations in more complex problems.

In summary, the problem of prediction, defined as exploiting correlations about the past dynamics to anticipate the future state comes up in many biological systems from motion prediction to evolution. This problem can be formulated in the same way, although as we have shown, the details of the dynamics matter for how best to encode a predictive representation and maximize the information the system can retain about the future state. Dynamics that results in Gaussian propagators is most informatively predicted using Gaussian representations. However non-Gaussian propagators introduce disjoint non-Gaussian representations that are nevertheless predictive.

By providing a set of dissected solutions to the predictive information bottleneck problem, we hope to show that not only is the approach feasible for biological encoding questions, it also illuminates connections between seemingly disparate systems (such as visual processing and the immune system). In these systems the overarching goal is the same, but the microscopic implementation might be very different. Commonalities in the optimally predictive solutions as well as the most generalizable ones can provide clues about how to best design experimental probes of this behavior, at both the molecular and cellular level or in networks.

## Acknowledgments

This work was supported in part by the US National Science Foundation, through the Center for the Physics of Biological Function (PHY–1734030), and a CAREER award to SEP (1652617); by the National Institutes of Health BRAIN initiative (R01EB026943–01); by a FACCTS grant from the France Chicago Center; and by European Research Council Consolidator Grant (724208).

## IV. SUPPORTING INFORMATION

## Appendix A COMPUTING THE OPTIMAL REPRESENTATION FOR JOINTLY GAUSSIAN PAST-FUTURE DISTRIBUTIONS

We follow Chechik, et al.[25] to analytically construct the optimally predictive representation variable, 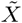, when the input and output variables are jointly Gaussian. The input is 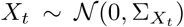 and the output is 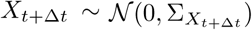. The joint distribution of *X*_*t*_ and *X*_*t*+∆*t*_ is Gaussian. To construct the representation, we take a noisy linear transformation of *X*_*t*_ to define 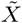

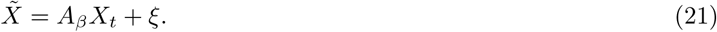

Here, *A*_*β*_ is a matrix whose elements are a function of *β*, the tradeoff parameter in the information bottleneck objective function between compressing, in our case, the past while retaining information about the future. *ξ* is a vector of dimension dim(*X*_*t*_). The entries of *ξ* are Gaussian-distributed random numbers with 0 mean and unit variance. Because the joint distribution of the past and the future is Gaussian, to capture the dependencies of *X*_*t*+∆*t*_ on *X*_*t*_ we can use a noisy linear transform of *X*_*t*_ to construct a representation variable that satisfies the information bottleneck objective function[25].

We compute *Aβ* by first computing the left eigenvectors and the eigenvalues of the regression matrix, 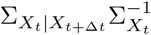. Here, 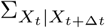 is the covariance matrix of the probability distribution of 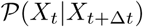. These eigenvector-eigenvalue pairs satisfy the following relation

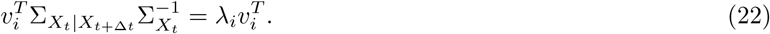

(We are taking 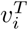 to be a row vector, rather than a column vector.)

The matrix, *A*_*β*_, is then given by

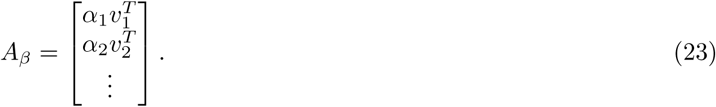

*β*_*i*_ are scalar values given by

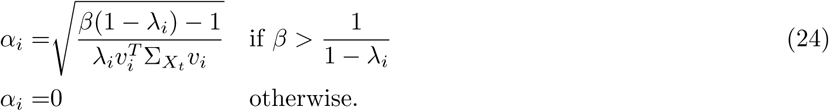

The α_*i*_ define the dimensionality of the most informative representation variable, 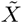. The dimension of 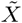 is the number of non-zero α_*i*_. The optimal dimension for a given *β* is, at most, equal to the dimension of *X*_*t*+∆*t*_. The set of values, 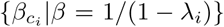, can be thought of as critical values, as each 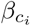 triggers the inclusion of the *i*th left eigenvector into the optimal *X*. The critical values depend strongly on the particular statistics of the input and output variable, so they may be different as the parameters that generate *X* change.

To compute the information about the past and future contained in 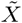, we compute 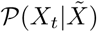 and 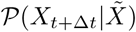. These distributions are Gaussian. The mean of each distribution corresponds to the encoded value of *X*_*t*_ and *X*_*t*+∆*t*_. The variance corresponds to the uncertainty, or entropy, in this estimate. To compute the variance, we need the variance of 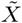

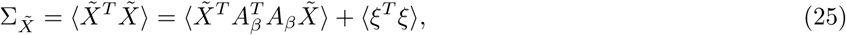

where the excluded terms are zero. Recalling the definition of *ξ*, we can simplify this expression to yield

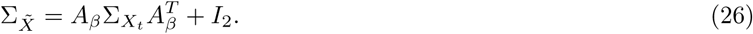

Here, *I*_2_ is the identity matrix. To compute the mutual information quantities, we use the following equations,

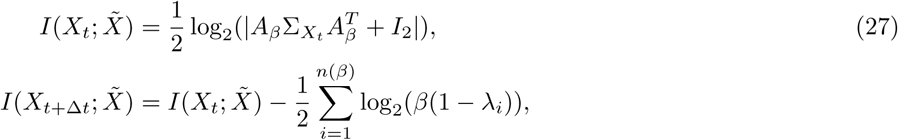

where *n*(*β*) corresponds to the number of dimensions included in *A*_*β*_. We also need the cross covariances between 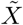 and *X*_*t*_ and between 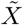 and *X*_*t*+∆*t*_, which are particularly useful for visualizing the optimal predictive encoding. To obtain these matrices, we use

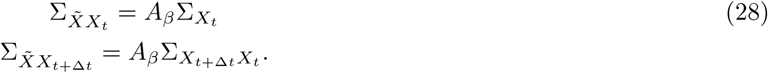

We can use these results and the Schur complement formula to obtain

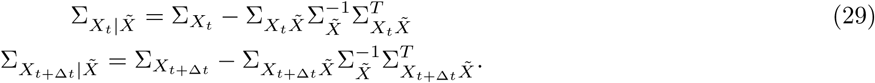

## Appendix B THE STOCHASTICALLY DRIVEN DAMPED HARMONIC OSCILLATOR

### .1 Harmonic Oscillator Model With No Memory

We begin by considering a mass attached to a spring undergoing viscous damping. The mass is being kicked by thermal noise. This mechanical system is largely called the stochastically driven damped harmonic oscillator (SDDHO). A simple model for its position and velocity evolution is given by

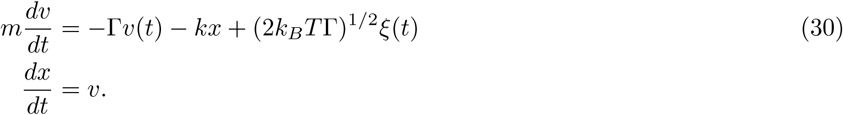

We use the redefined variables presented in the main text Equations 2 − 9 to rewrite the equations as

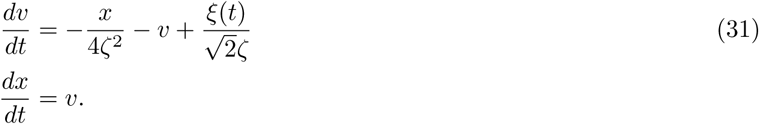

There are now two key parameters to explore: *ζ* and ∆*t*. There are three regimes of motion described by this model. The overdamped regime occurs when *ζ* > 1. In this regime of motion, the mass, when perturbed from its equilibrium position, relaxes back to its equilibrium position slowly. The underdamped regime occurs when *ζ* < 1. In this regime of motion, when the mass is perturbed from its equilibrium position, it oscillates about its equilibrium position with an exponentially decaying amplitude. At *ζ* = 1, we are in the critically damped regime of motion; in this regime, when the mass is perturbed from equilibrium, it returns to equilibrium position as quickly as possible without any oscillatory behavior.

To apply the information bottleneck method to this system, we need to compute the following covariance and cross covariance matrices: 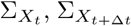, and 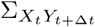. We note that because the defined motion model is stationary in time, 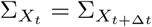. Using the procedure given in Flyvbjerg et. al. [52], we can compute the requisite autocorrelations to describe the cross-covariance matrix, 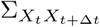.

We begin by using the equipartition theorem that states that

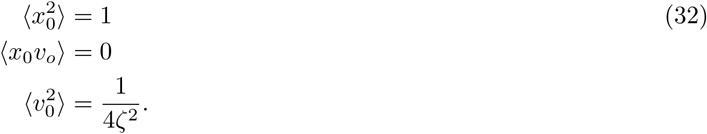

The covariance matrices are symmetric, so we can use these values to define the elements of 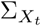. We then obtain expressions for 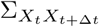

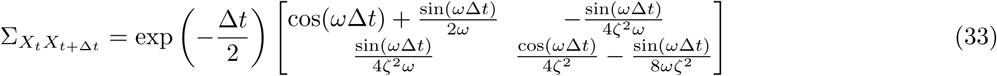

where we have defined 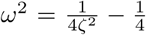. An alternative approach for the derivation of the above correlation values by methods of Laplace transforms can be found in Sandev et. al. [29].

To construct the optimal representation for prediction, we need the conditional covariance matrices, 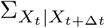 and 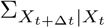. This can be computed using the Schur complement formula to yield

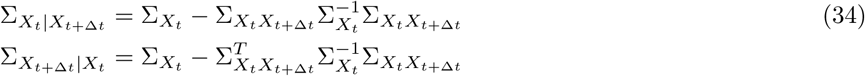

We provide a graphical representation of these distributions in Fig. 2b (main text). These graphical representations correspond to the contour inside which ~ 68% of observations are observed (i.e. one standard deviation from the mean).

### .2 Applying the information bottleneck Solution

To apply the information bottleneck solution, we construct the matrix, 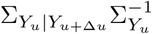, and find its eigenvalues and eigenvectors. The left eigenvectors of the matrix will be denoted by the columns of a new matrix, *w*, given by

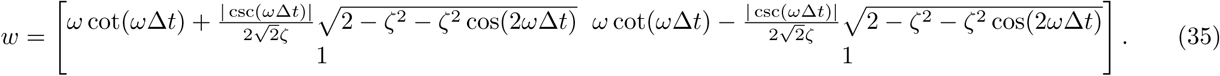

The eigenvalues are then

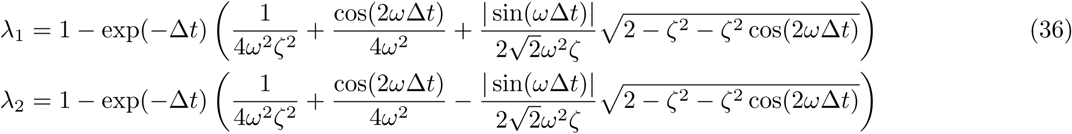

The transformation matrix, *A*_*β*_, will now depend on the parameters of the stimulus. Hence, we now refer to this matrix as *A*_*β*_ (*ζ*, ∆*t*), illustrating its functional dependence on those parameters.

Some general intuition can be gained from the form of the above expressions. The eigenvalue gap, λ_1_ − λ_2_ is proportional to 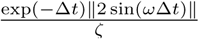. This suggests that the eigenvalue gap grows for small ∆*t*, then shrinks for large *∆t*. Additionally, in the small ∆*t* limit, the eigenvectors align strongly along the position and velocity axes, with the eigenvector corresponding to the smaller eigenvalue being along the position axis. Hence, for predictions with small *∆t*, the representation variable must encode a lot of information about the position dimension. For longer timescale predictions, both eigenvectors contribute to large levels of compression, suggesting that the encoding scheme should feature a mix of both position and velocity. This is presented in Figure 5.

We also compute the total amount of predictive information available in this stimulus. This is given by

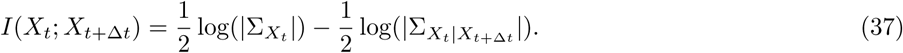

Simplifying this expression yields

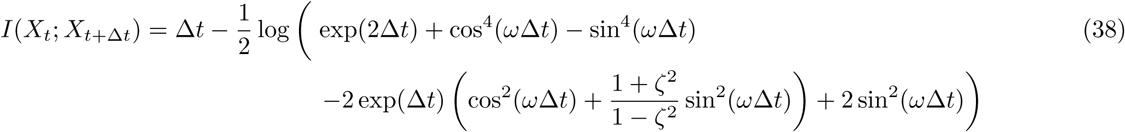

We can see for very large ∆*t*, this expression becomes

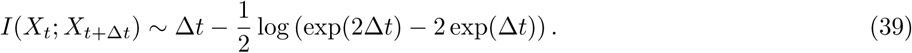

For small ∆*t*, we note there are two conditions: 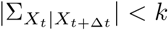 and 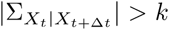, where *k* corresponds to width of the distribution. If the width of the Gaussian is below *k*, we treat this as being effectively deterministic. In this case,

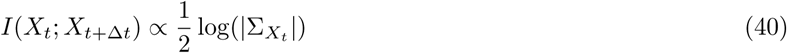

where there are some constants that set the units of the information and the reference point. For widths larger than

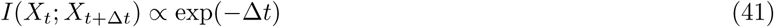

### .3 Comparing the information bottleneck Method to Different Encoding Schemes

We compare the encoding scheme discovered by the information bottleneck to alternate encoding schemes. We accomplish this by computing the optimal transformation for a particular parameter set for some value of β, *A*_*β*_ (*ζ*, ∆*u*). We then determine the conditional covariance matrix, 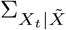. We generate data from this distribution and apply a two-dimensional unitary rotation. We then compute the covariance of the rotated data. This gives us a suboptimal encoding scheme, as represented in Figure 4b in yellow. We note that this representation contains the same amount of mutual information with the past as the optimal representation variable, though the dimensions the suboptimal encoding scheme emphasizes are very different. Evolving the rotated data forward in time and then taking the covariance of the resulting coordinate set gives us 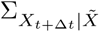, as plotted in Figure 4b in purple. We clearly see that encoding the past with the suboptimal representation reduces predictive information, as the predictions of the future are much more uncertain.

### .4 Comparing the information bottleneck method to Kalman filters

The Kalman filter approach seeks to fuse a prediction of a system’s coordinates at time *t* + ∆*t* based on initial coordinates at time *t* and knowledge of the dynamical system with an observation at time *t*+∆*t* to increase the certainty in the inference of the coordinates at time *t* + ∆*t*[26]. We present Kalman filters here to highlight the differences between the information bottleneck method and Kalman filtering techniques. First, we consider the structure of the Kalman filter,

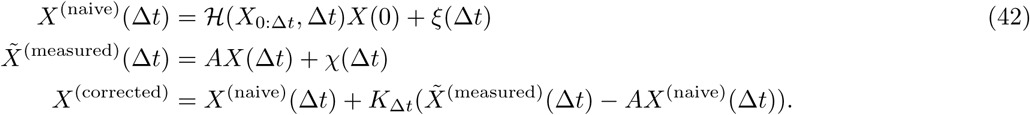

The first equation here considers an initial condition, *X*(0), a dynamical system model, 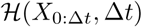, and a particular noise condition to construct an estimate of where an observer might expect their system to be after some time, ∆*t*, has passed from initial time 0. The second equation constructs a measurement, 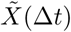(∆*t*) of the true coordinates, using a known observation model, *A*, and some measurement noise, *χ*(∆*t*). *A* is analogous to the probabilistic mapping constructed in the information bottleneck scheme, 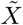; however, unlike in information bottleneck, in Kalman filtering, *A* is given to the algorithm and not discovered by any optimization procedure. Finally, the third equation unites the measurement, 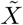, and the guess, *X*(∆*u*), by choosing *K*∆_*t*_, to be the transform which minimizes the variance between the true coordinates and the corrected coordinates. This correction is a post hoc correction and is not present in the information bottleneck scheme.

We now compare the results from a Kalman filtering technique and the information bottleneck when they both use the same probabilistic mapping, 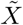. From Figure B.1(a), we see that even though the Kalman filter has higher levels of *I*_past_ for the same probabilistic mapping, the Kalman filtering algorithm is not efficient, as it is not extracting the most available predictive information.

### .5 An approach to encoding when the parameters of the stimulus are evolving

We examine prediction in the SDDHO when the underlying parameters governing the trajectory are evolving faster than adaptation timescales. While there are many possible strategies for prediction in this regime, we consider a strategy where the system picks a representation that provides a maximal amount of information across a large family of stimulus parameters. We chose this strategy because it enables us to analyze the transferability of representations from one parameter set against another. In other words, we can understand how robust representations learned for particular stimulus parameters are.

We first determine the predictive information extracted by an efficient coder for a particular representation level, *I*_past_ for a particular stimulus with parameters (*ζ*, ∆*t*), 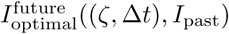. This predictive mapping is achieved by having a mapping, 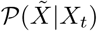. We apply this mapping to a new stimulus with different parameters (*ζ*, ∆*t*) to determine the amount of predictive information extracted by this mapping on a different stimulus with parameters (*ζ′*, ∆*t′*). We call this predictive information 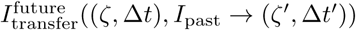.

We quantify the quality of these transferred representations in comparison with 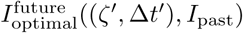 as

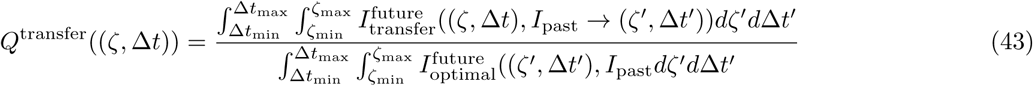

**FIG. B.1:**
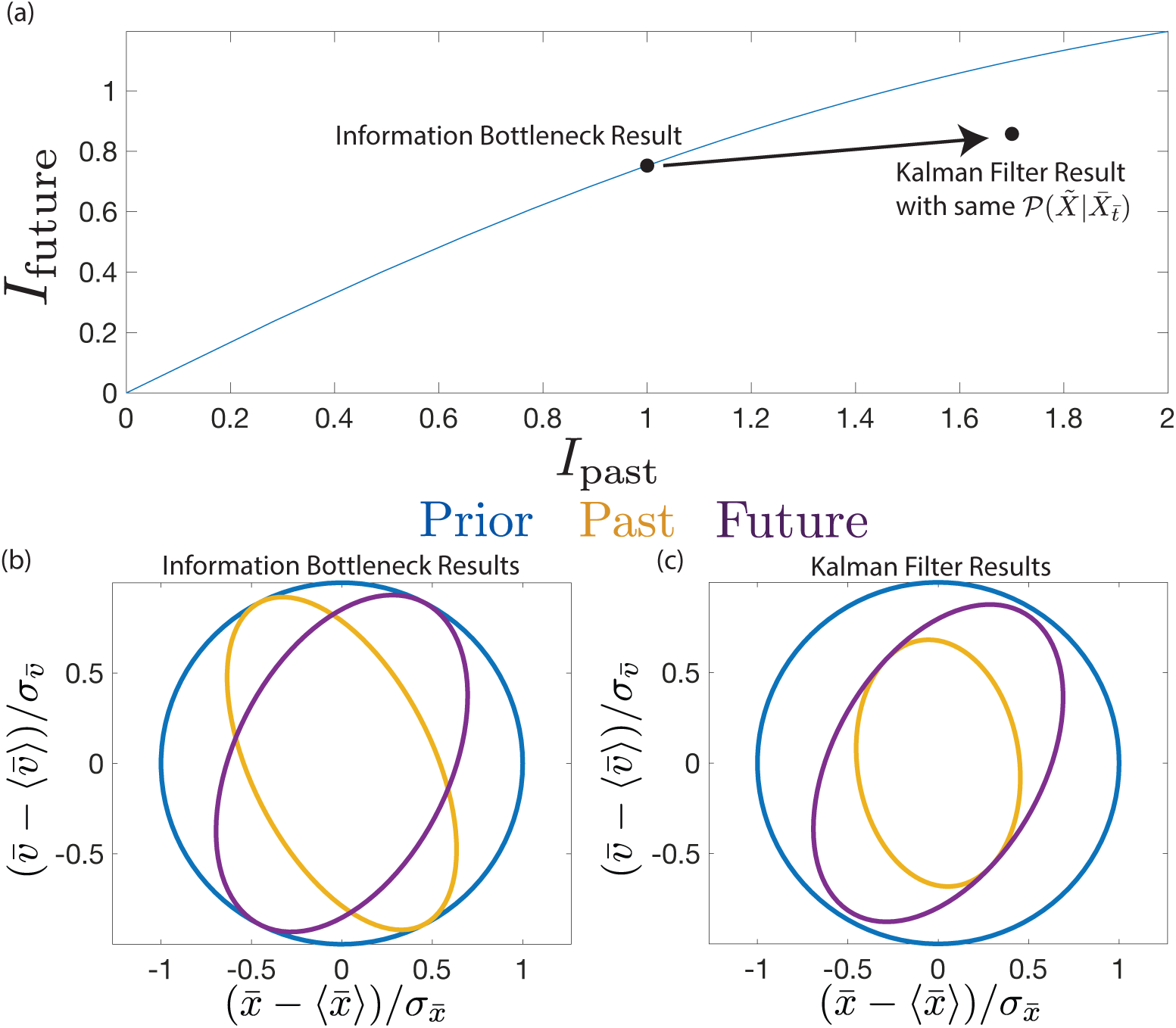
Kalman filtering schemes are not efficient coders for a given channel capacity. (a) Here, we present the information bottleneck curve for a stochastically driven damped harmonic oscillator with 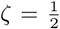 and ∆*t* = 1. We determine the optimal mapping for *I*_past_ = 1 bit and plot that point along the information bottleneck curve. Using the same probablistic mapping, we apply a Kalman filtering approach. We see that the Kalman filter approach results in an increase in both *I*_past_ and *I*_future_, but the result does not lie along the curve, indicating the scheme is not efficient. Panels (b) and (c) present the results in terms of uncertainty reduction in each scheme.

The resulting value is the performance of the mapping against a range of stimuli. In Figure 6, we analyzed the performance of mappings learned on 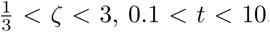, on stimuli with parameters 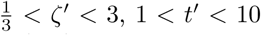. This choice of range is somewhat arbitrary, but it is large enough to see the asymptotic behavior in ∆*t*, *ζ*.

### .6 History Dependent Harmonic Oscillators

We extend the results on the Stochastically Driven Damped Harmonic Oscillator to history-dependent stimuli by modifying the original equations of motion to have a history dependent term using the Generalized Langevin Equation

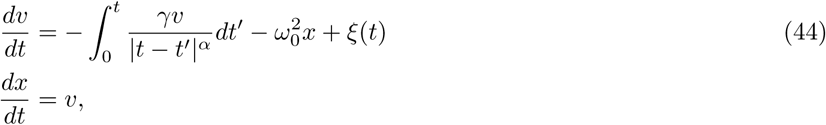

where 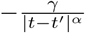 governs how the history impacts the velocity-position evolution. In the main text, we take *γ* = 1, *ω>* = 1, and *α* = 5/4. To compute the autocorrelation functions, we compute the Laplace transform of each autocorrelation function and numerically invert the Laplace transform to estimate the value

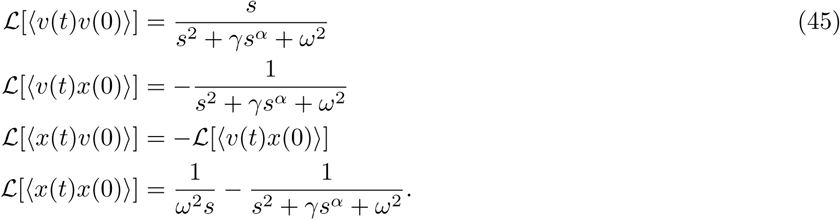

To expand our past and future variables to include multiple time points, we extend the past variable to be observations between *t − t*_0_ and *t* and the future variable to be *t* + ∆*t* to *t* + ∆*t* + *t*_0_. The size of the window is set by *t*_0_. We discretize each window with a spacing of *dt* = 2 and compute correlation functions along the discrete points of time, yielding the full covariance matrices. After this, the recipe is as outlined in Appendix A.

## Appendix C WRIGHT FISHER DYNAMICS

Wright-Fisher dynamics are used in population genetics to describe the evolution of a population of fixed size over generations. Here, we consider the diffusion approximation to the Wright-Fisher model with continuous time, given by Eq. 19. We numerically integrate Eq. 19 using a time step of *dt* = 0.001 and use 10000 data points starting from a given initial allele frequencies to estimate the joint distribution, *P* (*X*_*t*>+∆*t*_, *X*_*t*_). We discretize allele frequency space with *N* + 1 bins. We compute the maximum available predictive information for different values of the parameters (Fig. 10) using:

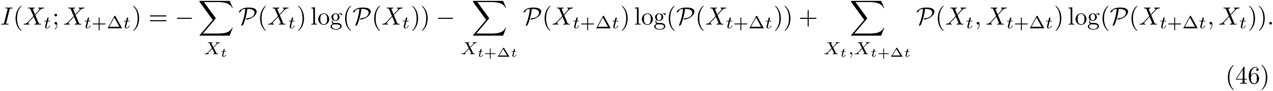

A simple estimate for 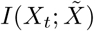 can be obtained by considering the case where each individual memory reflects a distinct cluster of allele frequencies. In the optimal encoding case, each memory encodes an equal amount of probability weight on the input variable[2, 53]. The upper bound on the information the representation variable has about the past state is 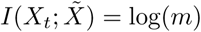.

**FIG. C.2:**
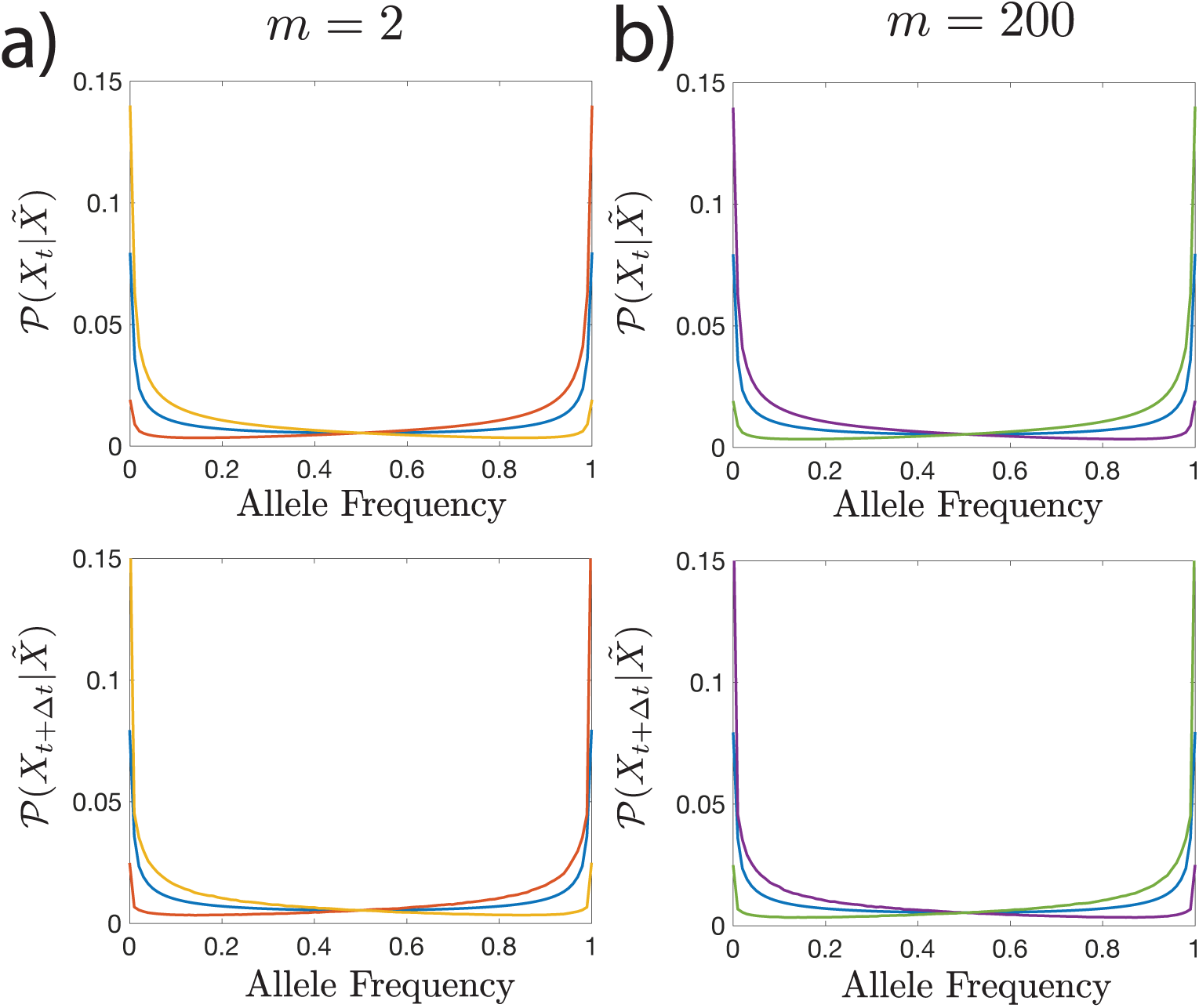
The optimal 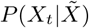 and 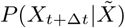 for Wright Fisher dynamics with *N* = 100, *N*_*µ*_ = 0.2, *N*_*s*_ = 0.001, ∆*t* = 1 with information bottleneck parameters *β* = 1.01 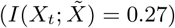 for *m* = 2 (a) and *m* = 200 (b). Many representations are degenerate in the *m* = 200 in this limit. The encoding schemes for *m* = 2 versus *m* = 200 are nearly identical for this small 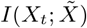) limit.

**FIG. C.3:**
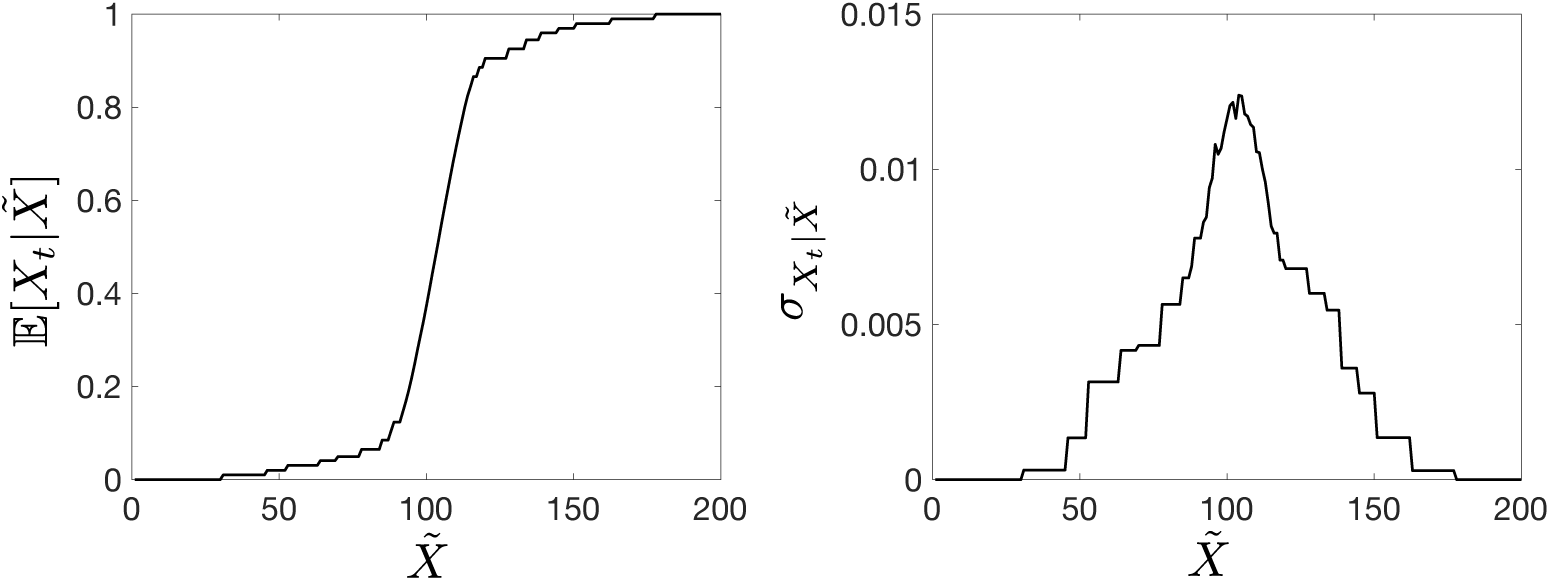
Mean (left) and variance (right) of the past allele frequency *X*_*t*_ conditioned on the (categorical) representation variable 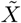 (left), for the information bottleneck solution of the Wright-Fisher dynamics with *m* = 200, *N* = 100, *N*_*µ*_ = 0.2, *N*_*s*_ = 0.001, *β* = ∞. The standard deviation is not constant: it is smaller where the prior probability of *X*_*t*_ is large.

